# Design and analysis of synthetic carbon fixation pathways based on novel enzymatic reactions

**DOI:** 10.1101/2025.11.21.689712

**Authors:** Vittorio Rainaldi, Sarah D’Adamo, Nico J. Claassens

## Abstract

Biological carbon fixation is currently limited to seven naturally occurring pathways, each with its own limitations and constraints. In recent years, computational analyses of known biochemical reaction networks have identified dozens of theoretical carbon fixation pathways, some of which may have the potential to outperform their natural counterparts. This mix-and-match approach, however, cannot account for those reactions that have not been reported to occur in nature, which heavily limits the possible solution space. Here, we use a bioretrosynthetic approach coupled with expert biochemical knowledge to identify several novel pathways that leverage enzyme promiscuity and the latent biochemical reaction space. We analyze the thermodynamic, stoichiometric, and kinetic parameters of these pathways and compare them to the ubiquitous Calvin-Benson-Bassham cycle and previously proposed synthetic CO_2_ fixation cycles, highlighting advantages and disadvantages. We identify several promising pathways that could potentially outcompete the Calvin cycle and other previously proposed synthetic CO_2_ fixation pathways in predicted biomass yield and/or overall pathway activity. In addition, unlike many of the previously proposed mix-and-match pathways, the pathways proposed in this work do not require vitamin B_12_, which is an advantage for future implementation in plants or microalgae that typically lack B_12_ biosynthesis. This work highlights the need for enzyme engineering and design in the quest for efficient biological carbon fixation.

## Introduction

The Calvin-Benson-Bassham (CBB) cycle is responsible for over 99% of net primary productivity on earth^1^. Despite its dominance, it is a relatively inefficient pathway, requiring a high energy investment in terms of ATP to fix carbon. Furthermore, its key enzyme, Rubisco, is rather slow when compared to other naturally occurring carboxylases and it presents a detrimental side activity with oxygen^2,3^, which results in additional energy costs and partial carbon loss during the cellular process known as photorespiration^4^.

Many synthetic carbon fixation pathways have been designed so far by using a mix- and- match approach that combine enzymes from disparate sources with fast and efficient carboxylases to generate theoretical cycles that have not been described in nature^5–7^. Nevertheless, a recent quantitative analysis of natural and synthetic carbon fixation pathways by L we and Kremling showed that most pathways are in fact inferior to the CBB cycle or only marginally more efficient in terms of biomass yield and pathway activity^7^, likely making the massive engineering efforts required to implement them unjustified.

Almost all of the current synthetic pathway designs, however, are limited by the fact that they are based only on known enzymatic activities from databases such as KEGG, which vastly limits the possible solution space. Erb, Jones, and Bar-Even have previously put forward a proposal for categorizing pathways based on five levels, with (1) naturally occurring pathways, (2) transplanted natural pathways into other organisms, (3) pathways designed by mix-and-match of only naturally existing enzymes (4) pathways including at least one novel enzyme reaction based on known enzymatic mechanisms and (5) pathways containing at least one new-to-nature enzymatic mechanism. Since designing enzymes with new-to-nature reaction mechanisms is still in its infancy despite recent progress^8,9^, we propose and analyze pathways in level 4. We examine three types of reactions: experimentally validated promiscuous activities, proposed promiscuous activities based on close structural similarity with the substrate of an existing enzyme, and proposed activities based on chemical similarity with other substrates of existing enzymes (see Methods).

In this study, we use a combination of bioretrosynthesis and expert biochemical knowledge to generate novel pathway designs and analyze their thermodynamic^10^, stoichiometric^11^, and kinetic^12^ parameters, comparing them to the CBB cycle and highlighting advantages and disadvantages. All these pathways require engineering of at least one enzyme to change its substrate or cofactor specificity or other relevant parameters such as oxygen tolerance. Oxygen tolerance is particularly important in the context of engineering CO_2_ fixation in plants and algae, since these organisms produce oxygen as part of photosynthesis, which would lead to the rapid inactivation of oxygen-sensitive enzymes. However, oxygen tolerance is also important in hydrogen-based microbial bioproduction from CO_2_, as CO_2_ fixation and bioproduction pathways usually require ATP, which is typically generated from O_2_-dependent respiration, and alternative electron acceptors (e.g. nitrate, sulphate) are not sufficiently scalable for large-scale biotechnological production. Realizing the proposed pathways will require varying degrees of enzyme engineering efforts and is beyond the scope of this work. However, we believe that given the recently demonstrated examples of engineered enzymes, as well as advances in computational approaches for enzyme engineering and design, many of these designs may become feasible in the near future.

Overall, this paper provides pathway designs based on novel enzymes and highlights the need for enzyme engineering to achieve efficient carbon fixation, while offering suggestions regarding possible starting points to engineer the proposed pathways.

## Results

### The β-glutamate cycle

We recently proposed and showed the partial in vivo implementation of a new family of carbon fixation pathways called the reductive methylaspartate (rMASP) cycles using the standard mix-and-match approach^13^. While these cycles are promising, they suffer from two main issues. First, the enzyme crotonyl-CoA carboxylase/reductase (CCR) is relatively inefficient because the NADPH invested for its reductive carboxylation activity is later only partially recovered in the form of a reduced quinone, leading to a loss of energy (3 protons or roughly 1 ATP). Second, these cycles require vitamin B_12_ as a cofactor for two enzymatic reactions. Vitamin B_12_ is a highly complex cofactor, which cannot be synthesized by most organisms, importantly including all higher plants and many algae^14^, thus limiting the applicability of pathways containing B_12_-dependent enzymes mostly to bacteria.

To overcome both issues, we designed a variant of the rMASP cycles that we name the β-glutamate cycle based on its characteristic metabolite (Fig. 1). According to our computational analysis, the β-glutamate cycle shows favorable parameters when compared to the CBB cycle, including higher max-min (thermodynamic) driving force (MDF), higher theoretical biomass yield, and comparable pathway activity. These advantages increase in ambient CO_2_ environments, with a ∼40% increase in yield (assuming 20% Rubisco oxygenation activity for the CBB cycle at ambient CO_2_^15–17)^ (Table S1).

**Figure 1:**
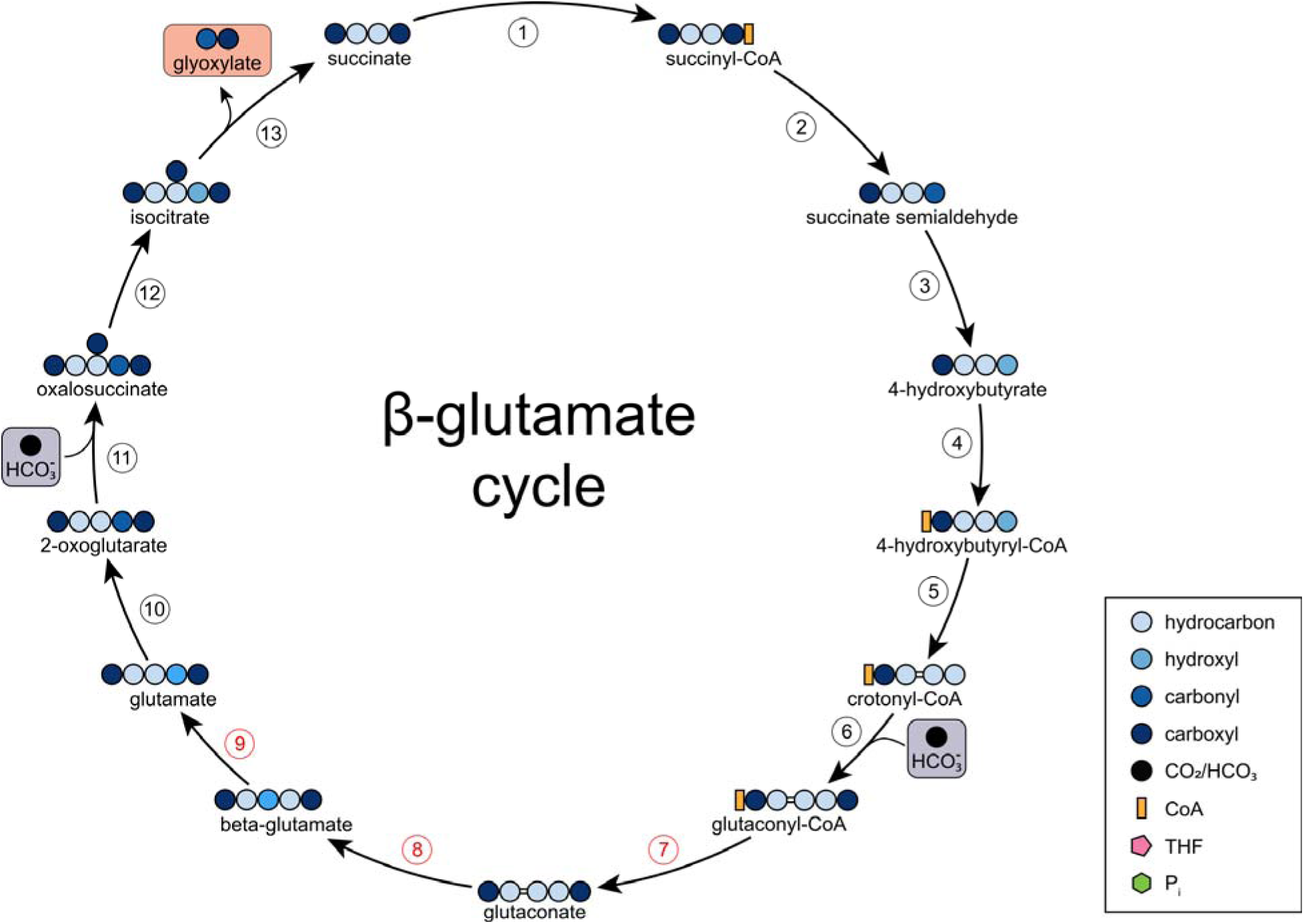
The β-glutamate cycle. In this figure and following ones, colored spheres represent different carbon groups and colored shapes represent cofactors or other molecules as represented in the legend. Both input and output molecules are framed by boxes. 1, succinyl-CoA synthetase; 2, succinyl-CoA reductase; 3, succinate semialdehyde dehydrogenase; 4, 4-hydroxybutyrate CoA transferase; 5, 4-hydroxybutyryl-CoA dehydratase; 6, crotonyl-CoA carboxylase; 7, glutaconate CoA transferase; 8, β-glutamate ammonia lyase; 9, glutamate 2,3-aminomutase; 10, glutamate dehydrogenase; 11, 2-oxoglutarate carboxylase; 12, isocitrate dehydrogenase; 13, isocitrate lyase. In this figure and following ones the enzymes marked in red have never been reported in the literature. Bioprospecting or engineering efforts will likely be required to establish their activities.

The β-glutamate cycle does not use CCR but instead depends on the promiscuous activity of methylcrotonyl-CoA gamma carboxylase (MCC)^18,19^, which we will refer to as crotonyl-CoA gamma carboxylase (CCC). This biotin-dependent carboxylase is involved in leucine degradation across diverse phyla including bacteria^20,21^ and humans^22^. Its promiscuous side activity on the slightly smaller crotonyl-CoA has been reported in the literature with about 90% of the reaction rate when compared to the native substrate. Since the carboxylation happens in gamma, as opposed to alpha, it produces glutaconyl-CoA instead of ethylmalonyl-CoA, thus obviating the need for a B_12_ dependent mutase to “straighten” the carbon skeleton.

The cycle proceeds with the removal of the CoA from glutaconyl-CoA via a CoA transferase or CoA ligase to yield glutaconate, which is further aminated to 3-aminoglutarate, also known as β-glutamate, which gives its name to the cycle. This amination reaction could be promiscuously performed by aspartate/methylaspartate ammonia lyase^23^, though most likely it would require engineering to expand the substrate binding pocket to accommodate one extra carbon. An alternative to this route could be the direct amination of glutaconyl-CoA, for example by a promiscuous β-alanyl-CoA ammonia lyase, an enzyme found in anaerobic bacteria and used for β-alanine fermentation^24^. The resulting β-glutamyl-CoA would once again require a CoA transferase or CoA ligase to remove the CoA and yield β-glutamate.

At this point, β-glutamate must undergo isomerization to produce glutamate. This reaction is catalyzed by the naturally occurring enzyme glutamate 2,3-aminomutase^25^, which is present in bacteria and archaea, respectively for glutamate fermentation and for the production of β-glutamate as an osmolyte^26^. This reaction, however, is highly challenging, requiring a radical mechanism based on S-adenosyl-methionine (SAM), making the enzyme oxygen-sensitive. Nevertheless, there are oxygen-tolerant variants reported to catalyze the same reaction aerobically, though using lysine as a substrate as opposed to glutamate^27–29^. Not much is known about this enzyme class, so it is difficult to know whether it would be easier to engineer oxygen-tolerance in a glutamate aminomutase or change substrate specificity in a lysine aminomutase. It must be noted, however, that while there are many examples of oxygen-tolerant radical SAM enzymes^30,31^, e.g. *bioB* and *lipA* from *E. coli*, there seems to be a tradeoff between activity and oxygen tolerance, with oxygen-sensitive variants showing higher activity. For this reason, it may not be possible to obtain an oxygen-tolerant aminomutase with comparable kinetic parameters to oxygen-sensitive isoforms.

### The 3-hydroxyglutaryl-CoA lyase-methylcitrate (3HGL-MC) cycle

We reasoned that the product of CCC, glutaconyl-CoA, could also undergo hydration rather than amination, yielding 3-hydroxyglutaryl-CoA. This promiscuous reaction is indeed reported to occur efficiently^32^. This metabolite could then be split by 3HMG-CoA lyase to acetyl-CoA and malonate semialdehyde, with the latter being reduced to propionyl-CoA and subsequently going through the methylcitrate cycle for propionyl-CoA assimilation^33,34^. The resulting succinate can then be activated to succinyl-CoA and reduced to crotonyl-CoA to restart the cycle (Fig. 2). More pathways based on CCC and glutaconyl-CoA hydration have previously been reported^5,35^, however the authors describe a hydration in C2 rather than C3, which produces 2-hydroxyglutaryl-CoA. Unfortunately, the enzyme that catalyzes this reaction is based on a ketyl radical mechanism and hence extremely oxygen-sensitive^36,37^. To the best of our knowledge there has never been a report of ketyl radical enzymes operating in aerobic conditions, making these proposed routes infeasible for aerobic implementation, while their anaerobic implementation would require an external electron acceptor other than oxygen for respiration, making the pathways not relevant for implementation in algae or plants and most other biotechnological applications. The 3HGL-MC cycle, on the other hand, does not use any oxygen-sensitive enzymes, since hydration in C3 occurs easily due to the differences in the pKa between the C2 protons and the C3 protons^38^.

**Figure 2:**
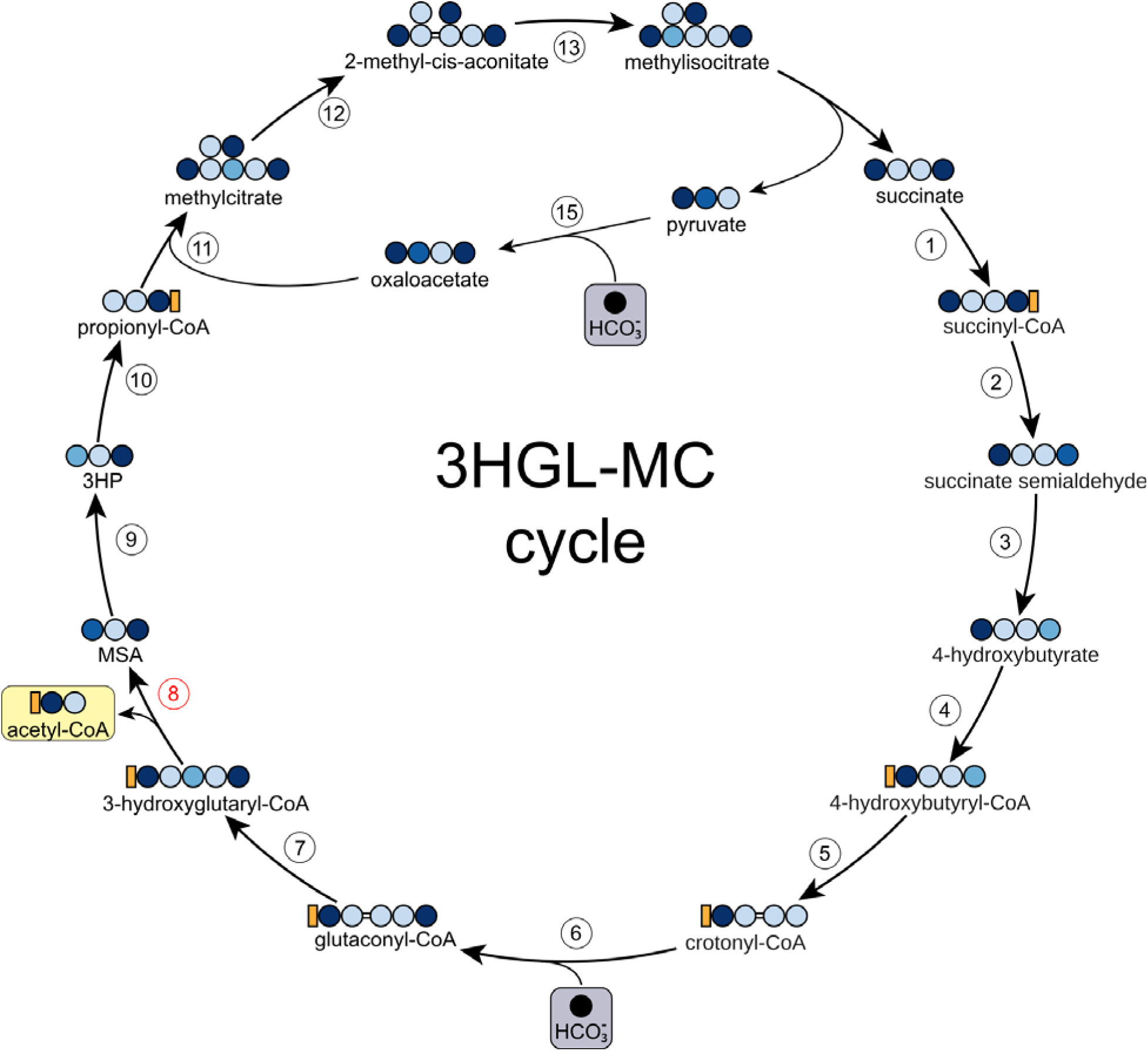
The 3-hydroxyglutaryl-CoA lyase-methylcitrate cycle. 1, methylmalonyl-CoA mutase; 2-6 are as in figure 1; 7, glutaconyl-CoA hydratase; 8, 3-hydroxyglutaryl-CoA lyase; 9, malonate semialdehyde reductase; 10, propionyl-CoA synthase; 11, methylcitrate synthase; 12, methylcitrate dehydratase; 13, aconitase; 14, methylisocitrate lyase; 15, pyruvate carboxylase.

This pathway is predicted to have about 25% higher biomass yield at ambient CO_2_, with roughly double the activity (Table S1, Fig. 6). Moreover, the addition of acetyl-CoA carboxylase and malonyl-CoA reductase would make the cycle autocatalytic, much like the CBB cycle and reverse tricarboxylic acid (rTCA) cycle. Autocatalytic cycles are expected to be more stable and resilient to metabolic perturbations^39^, potentially simplifying engineering efforts. Two other variants of this pathway, the 3HGL-PCC cycle (Fig. S1) and the 3HGM cycle (Fig. S2) can be envisioned, however their strict B_12_ requirement limits their potential application to some bacteria and algae^14^.

### The α-ketobutyrate carboxylase (ABC) cycles

While the pathways based on crotonyl-CoA carboxylation discussed above use either the condensation of two acetyl-CoA molecules or the reduction of succinyl-CoA to form a C4 metabolite for carboxylation, a different strategy can be envisioned based on the threonine biosynthesis pathway. The in vivo implementation of the Serine Threonine Cycle (STC) for full formatotrophic growth was recently reported in *E. coli*^40^.

The STC relies on the condensation of glycine derived from threonine cleavage with externally supplied formate and an additional carboxylation reaction based on pyruvate carboxylase to generate the C4 molecule aspartate, which is the precursor for threonine. To convert this formate assimilation pathway into a CO_2_ fixation pathway design, we reasoned that threonine could instead undergo deamination to produce α-ketobutyrate. The latter C4 metabolite is an α-keto acid highly similar to pyruvate and thus amenable to carboxylation via an engineered pyruvate carboxylase. While we could find no reports of a biotin-dependent α-ketobutyrate carboxylase in the literature, there is at least one study showing partial promiscuous activity of pyruvate carboxylase on α-ketobutyrate^41^. This limited activity could be used as the basis for mutagenesis or rational engineering in an approach similar to the one used to change the substrate specificity of propionyl-CoA carboxylase to glycolyl-CoA^42^. An α-ketobutyrate carboxylase (ABC) would form the basis of the ABC cycles we discuss in this work.

While pyruvate carboxylation produces oxaloacetate, α-ketobutyrate carboxylation would produce 3-methyloxaloacetate (also known as 2-methyl-3-oxosuccinate or oxalopropionate). This compound can undergo reductive amination or transamination to methylaspartate^43^, reduction to 3-methylmalate (erythro or threo), or condensation with acetyl-CoA to methylcitrate, though we did not investigate the latter option further, since the resulting cycle is rather complex and inefficient (Fig. S3).

In the first case, methylaspartate can be isomerized to glutamate via a B_12_-dependent glutamate mutase (GM) and then proceed via the same steps as the previously described rMASP cycles^14^. The ABC-GM cycle (Fig. S4) compares favorably to the CBB at ambient CO_2_ in terms of biomass yield, though its strict requirement for B_12_ limits its applicability to microorganisms.

To bypass the B_12_ requirement, we propose a cycle where methylaspartate is deaminated to mesaconate and hydrated to citramalate, which is then activated to citramalyl-CoA and split to pyruvate and acetyl-CoA via a citramalyl-CoA lyase (CCL). The resulting ABC-CCL cycle (Fig. 3) compares favorably to the CBB cycle in terms of theoretical biomass yield and predicted pathway activity at ambient CO_2_, is insensitive to oxygen and does not require B_12_ as a cofactor (Table S1, Fig. 6). Conversion of methyloxaloacetate to mesaconate could proceed via different routes with slightly different energetic demands (Fig. S5).

**Figure 3:**
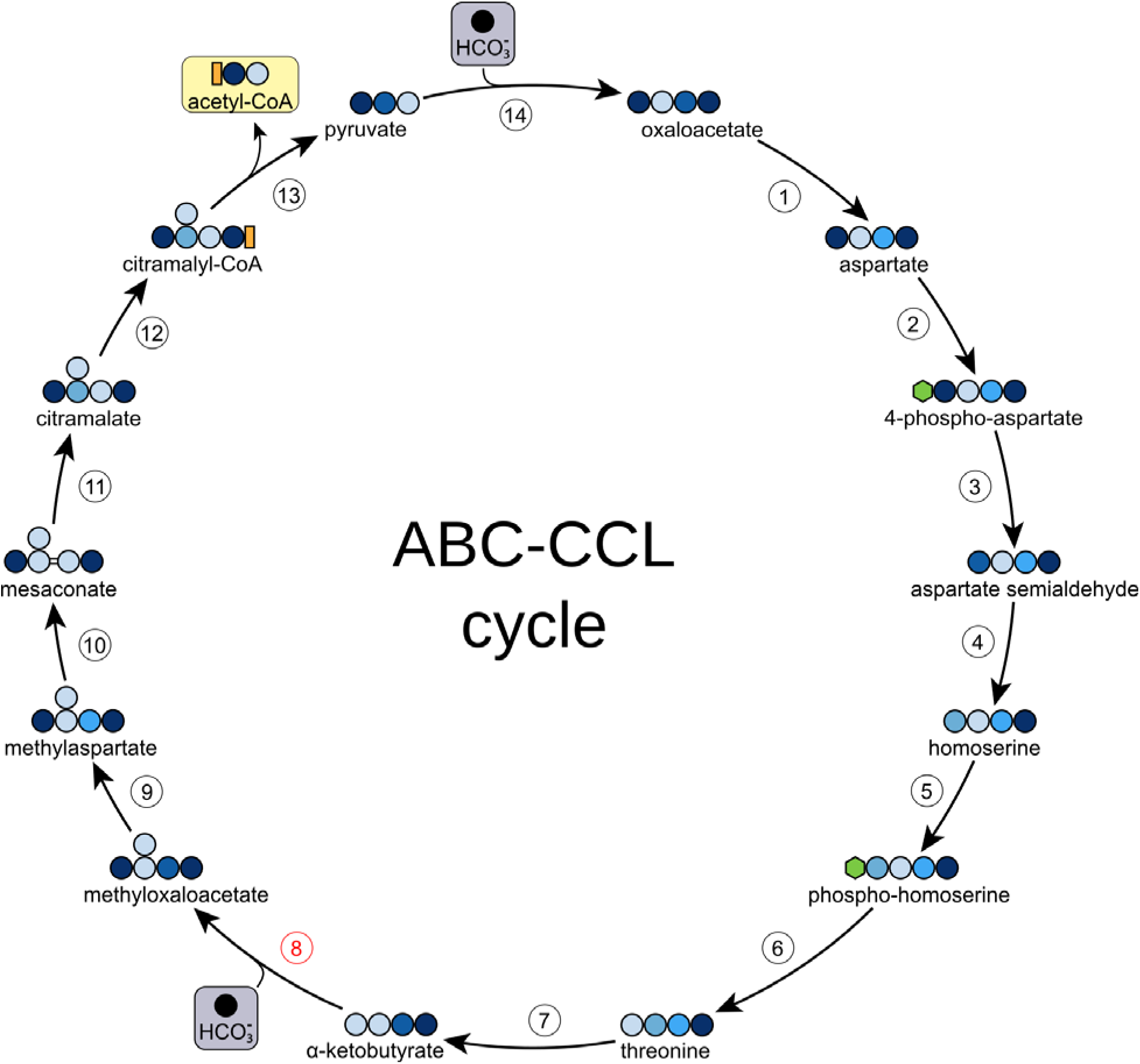
The α-ketobutyrate carboxylase-citramalyl-CoA lyase cycle. 1-9 are as in figure 4. 10, methylaspartate ammonia lyase; 11, mesaconase; 12, citramalate CoA transferase; 13, citramalyl-CoA lyase; 14, pyruvate carboxylase.

A third option could use the promiscuous activity of malate dehydrogenase (MDH) for the reduction of methyloxaloacetate to erythro-methylmalate, which can further be activated to a CoA ester and split to propionyl-CoA and glyoxylate. This variant, known as the ABC-PCC cycle (Fig. S6) also requires B_12_ for the conversion of methylmalonyl-CoA to succinyl-CoA following the carboxylation of propionyl-CoA.

Notably, all three proposed ABC cycles only require one enzyme to be engineered, and their overlap with the STC shows that they could be engineered relatively easily by applying some of the lessons learned during the STC engineering and evolution in *E. coli*^40^. Nevertheless, our theoretical analysis shows that threonine biosynthesis may limit the overall activity of these pathways, especially through the bottleneck enzyme homoserine dehydrogenase (Fig. S7). It is unclear whether faster variants of this enzyme can be found or engineered, and if the thermodynamic constraints of this biosynthetic route can be overcome, e.g. by optimizing enzymes and/or metabolite concentration. More research will be needed to ascertain whether this theoretical prediction will be verified experimentally.

### The Ribulose-5-phosphate carboxylase-Entner-Doudoroff (RED) cycle

While all the cycles presented so far have some advantages over the CBB, it should be noted that that they are all based on completely different architectures, thus requiring extensive engineering efforts to establish in organisms that already use the CBB cycle, including algae and plants. For this reason, we set out to establish a carbon fixation cycle that is similar in architecture to the CBB cycle but presents more favorable characteristics in terms of biomass yield and pathway kinetics. The GND-Entner-Doudoroff (GED) cycle has been previously proposed as a promising alternative to the CBB based on natural enzymes^44^, however it is severely limited by the low activity and affinity for CO_2_ of its signature enzyme, 6-phosphogluconate dehydrogenase (GND). This enzyme normally catalyzes the oxidative decarboxylation of 6-phosphogluconate, coupling the thermodynamically highly favorable decarboxylation reaction to the unfavorable reduction of NADP. This same mechanism can be found in at least two other central metabolic enzymes, namely the malic enzyme, which produces pyruvate and NADPH from malate and NADP, and isocitrate dehydrogenase, which produces α-ketoglutarate and NADPH from isocitrate and NADP.

Notably, in both of these cases there is a biotin-dependent carboxylase to overcome the unfavorable thermodynamics and kinetics of these reactions in the direction of carboxylation, which is then coupled to a dehydrogenase/reductase. Pyruvate is carboxylated to oxaloacetate, which is then reduced to malate, while α-ketoglutarate is carboxylated to oxalosuccinate and then reduced to isocitrate^45–47^. A similar idea could be applied to the product of GND, ribulose-5-phosphate. Here, we propose the RED cycle (Fig. 4), which is based on a mechanistically plausible biotin-dependent ribulose-5-phosphate carboxylase. This enzyme would generate 3-keto-6-phosphogluconate, which could then be reduced to 6-phosphogluconate. For this latter reaction GND itself could be used, provided the residues involved in the decarboxylation half-reaction are inactivated.

**Figure 4:**
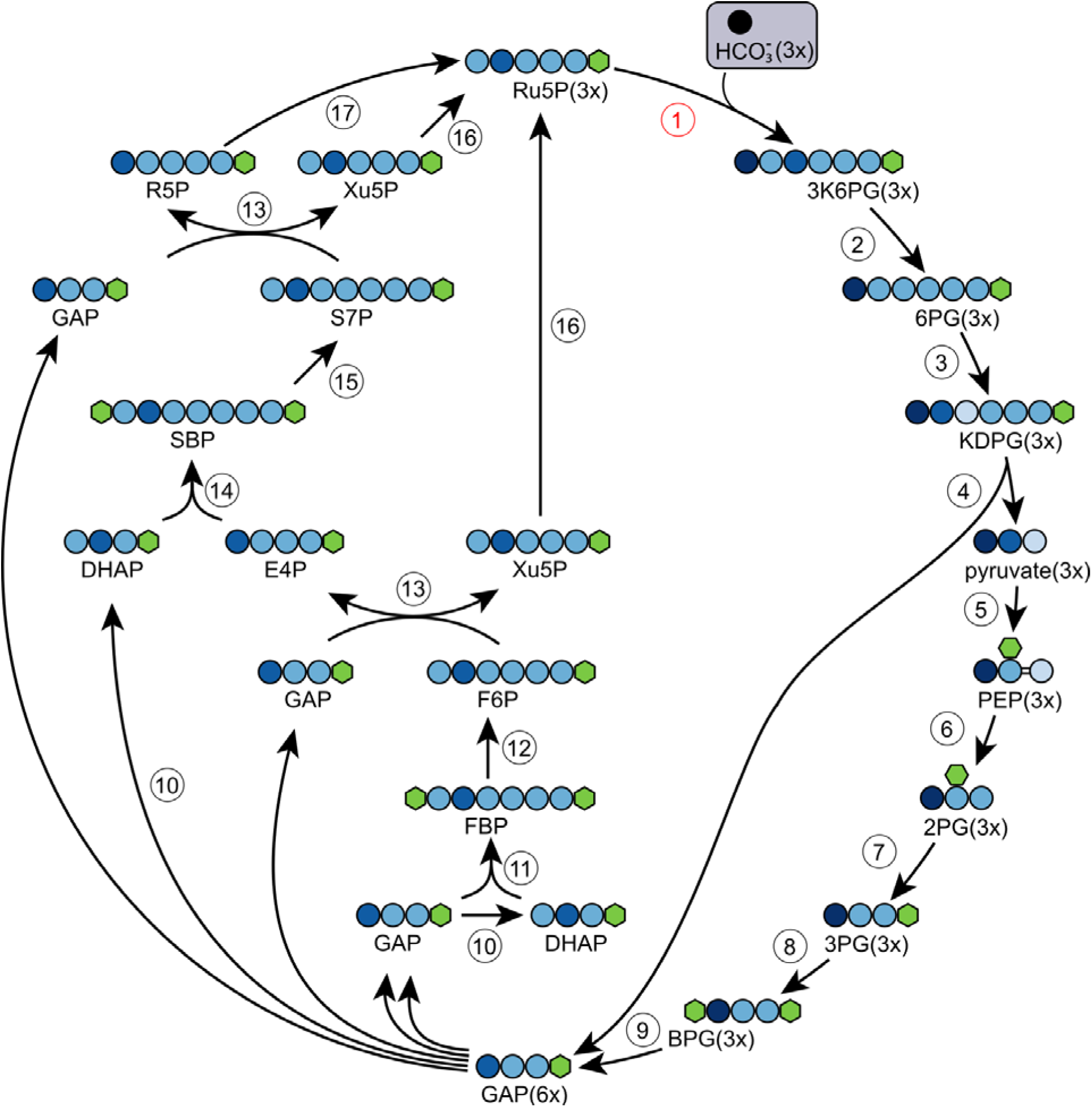
The Ribulose-5-phosphate carboxylase-Entner-Doudoroff (RED) cycle. 1, ribulose-5-phosphate carboxylase; 2, 6-phosphogluconate dehydrogenase; 3, Entner-Doudoroff dehydratase; 4, Entner-Doudoroff aldolase; 5, PEP synthetase; 6, enolase; 7, phosphoglycerate mutase, 8, phosphoglycerate kinase; 9, glyceraldehyde-3-phosphate dehydrogenase; 10, triose phosphate isomerase; 11, fructose bisphosphate aldolase; 12, fructose bisphosphatase; 13, transketolase; 14, sedoheptulose bisphosphate aldolase; 15, sedoheptulose bisphosphatase; 16, ribulose-5-phosphate epimerase; 17, ribulose-5-phosphate isomerase. 2PG, 2-phosphoglycerate; 3K6PG, 3-keto-6-phosphogluconate; 3PG, 3-phosphoglycerate; 6PG, 6-phosphogluconate; BPG, 1,3-bisphosphoglycerate; DHAP, dihydroxyacetone phosphate; E4P, erythrose-4-phosphate; F6P, fructose-6-phosphate; FBP, fructose-1,6-bisphosphate; GAP, glyceraldehyde-3-phosphate; KDPG, 2-keto-3-deoxy-phosphogluconate, PEP, phosphoenolpyruvate; R5P, ribose-5-phosphate; Ru5P, ribulose-5-phosphate; S7P, sedoheptulose-7-phosphate; SBP, sedoheptulose-1,7-bisphosphate; Xu5P, xylulose-5-phosphate.

To the best of our knowledge, there is no reported enzyme taking a similar substrate for the carboxylation reaction, meaning that an extensive engineering campaign will be necessary to change the substrate specificity. This is probably beyond what is achievable with current strategies based on mutagenesis and selection, however it may become possible in the future to apply novel computational enzyme design approaches to redesign the substrate pocket. The RED cycle is insensitive to the presence of oxygen, does not require CoA or exotic cofactors, it is favorable in terms of efficiency and kinetics at ambient CO_2_ (Table S1, Fig. 6), and it has a high degree of overlap with the CBB cycle, making it a good candidate for implementation in algae and plants. However, it should be noted that these organisms do not express lower glycolysis enzymes in their chloroplasts, which could increase the engineering complexity.

### The Dehydratase-less Ribulose-5-phosphate carboxylase-Entner-Doudoroff (DRED) Cycle

While the RED cycle is a promising alternative to the CBB cycle in terms of pathway activity, its ATP consumption is higher, and consequently its overall predicted biomass yield is lower under high CO_2_ conditions (Table S1, Fig. 6). This is due to the costly pyruvate kinase reaction, which consumes two ATP equivalents per pyruvate. To solve this issue, we propose a variant of the RED cycle that we name the DRED cycle (Fig. 5). This variant does not rely on the Entner-Doudoroff dehydratase (Edd) but instead uses an isomerase to convert 3-keto-6-phosphogluconate to 2-keto-6-phosphogluconate. This metabolite could subsequently serve as the substrate for the Entner-Doudoroff aldolase (Eda), producing glyceraldehyde-3-phosphate (GAP) and hydroxypyruvate. The latter can be reduced to glycerate, which is then activated to 2-phosphoglycerate and enters gluconeogenesis to generate another GAP. The advantage of this sequence of reactions is that glycerate kinase only consumes one ATP equivalent. As a result, this cycle is predicted have comparable yield and pathway activity to the CBB cycle even at high CO_2_ concentrations (Table S1, Fig. 6), all while eschewing the oxygenation side reaction of Rubisco.

**Figure 5:**
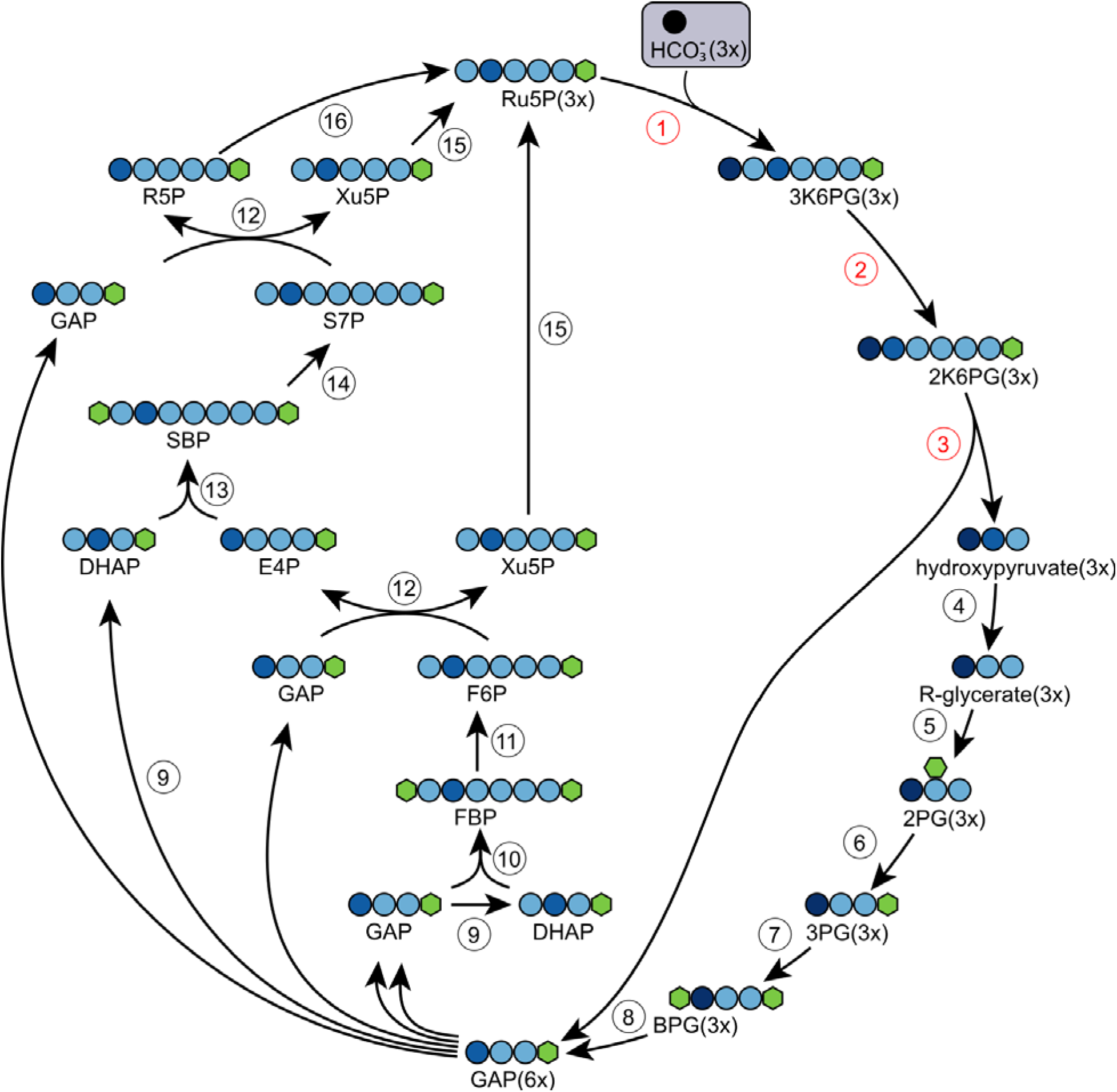
The Dehydratase-less-Ribulose-5-phosphate carboxylase-Entner-Doudoroff (DRED) cycle. 1, Ribulose-5-phosphate carboxylase; 2, 3-keto-6-phosphogluconate isomerase; 3, 2-keto-6-phosphogluconate aldolase; 4, hydroxypyruvate reductase; 5, glycerate kinase; 6, phosphoglycerate mutase, 7, phosphoglycerate kinase; 8, glyceraldehyde-3-phosphate dehydrogenase; 9, triose phosphate isomerase; 10, fructose bisphosphate aldolase; 11, fructose bisphosphatase; 12, transketolase; 13, sedoheptulose bisphosphate aldolase; 14, sedoheptulose bisphosphatase; 15, ribulose-5-phosphate epimerase; 16, ribulose-5-phosphate isomerase. 2K6PG, 2-keto-6-phosphogluconate; All other abbreviations are as in figure 5.

**Figure 6:**
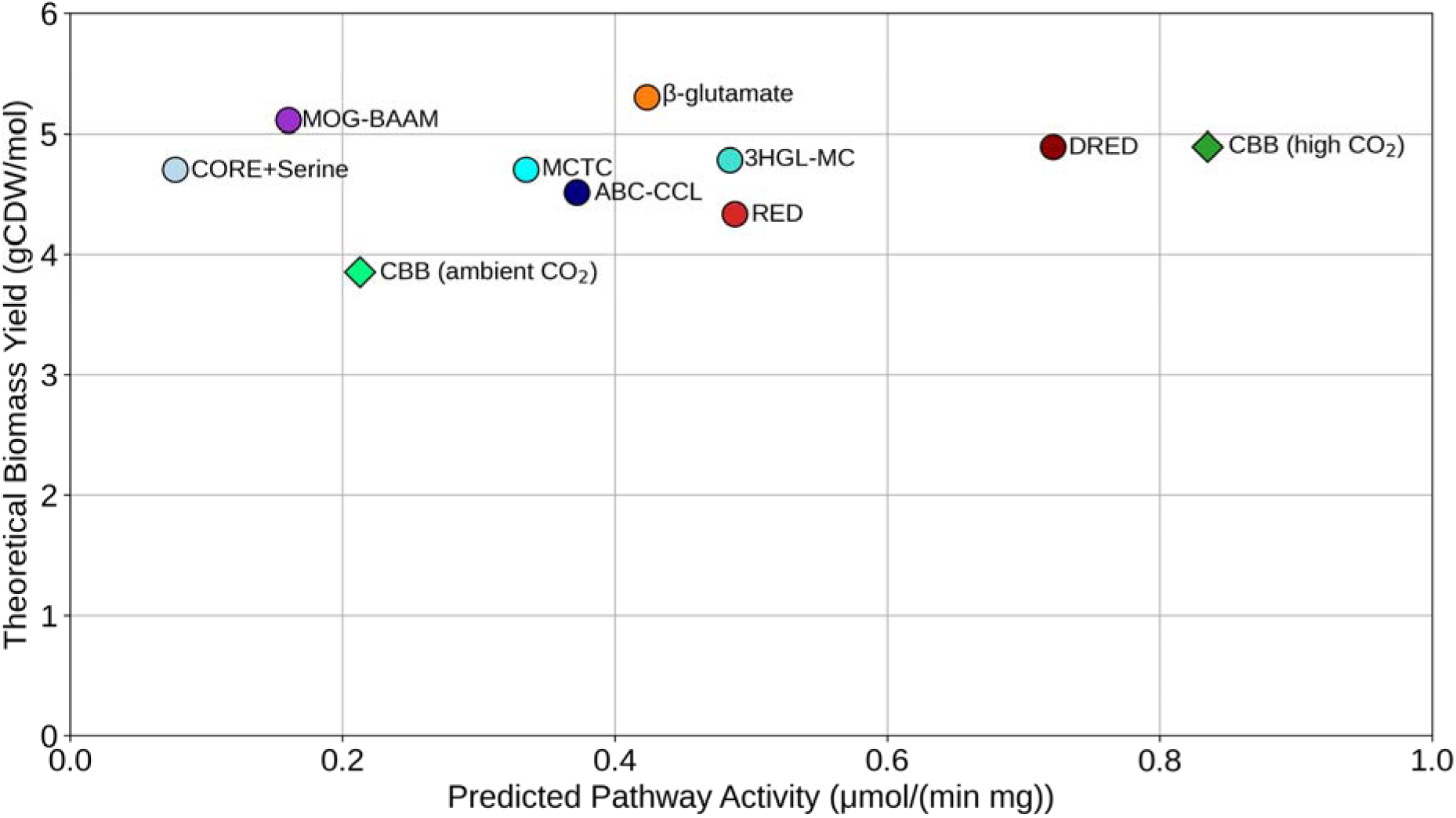
Plot of theoretical biomass yield vs predicted pathway activity for each pathway presented in this paper and previously proposed pathways. All pathways except the CBB cycle were modeled using ambient CO_2_ concentrations, see Methods. This figure includes only pathways that require enzyme engineering for their implementation (other than the CBB cycle). We also include comparisons to the previously proposed malyl-CoA-tartronyl-CoA cycle, also known as LATCH^51^ (Fig. S10) and the beta alanine variant of the malonyl-CoA-oxaloacetate-glyoxylate (MOG) cycle^52^ (Fig. S11). For additional comparisons to B12-dependent pathways proposed in this paper and previously proposed pathways, including the lactyl-CoA mutase variant of the MOG cycle^52,53^ (Fig. S12) and several mix-and-match CO_2_ fixation pathways^6,44,54–56^, see figure S13. A sensitivity analysis for all ECM values can be found alongside the ECM results in the online repository and a summary is available in Table S3.

Despite these advantages, the cycle requires the engineering of three enzymes, as opposed to only one for the RED cycle. Additionally, hydroxypyruvate is a moderately toxic metabolite which can inhibit bacterial growth at mM concentrations. While its effect on plants is not known, careful balancing of pathway enzymes is likely sufficient to keep the levels of hydroxypyruvate low and avoid its toxicity.

A similar way to bypass pyruvate kinase was proposed in the context of the GED cycle by Arren Bar-Even^48^, albeit using a 6-phosphogluconate-2-dehydrogenase (EC 1.1.1.43) instead of an isomerase, which is also a possibility here (Fig. S8).

### Coupling the CO_2_ reduction (CORE) cycle to formatotrophic pathways

An interesting alternative to CO_2_ fixation cycles was recently proposed by Satanowski, Marchal et al., who theorized several oxygen tolerant CO_2_ reduction (CORE) cycles and implemented one of them in *E. coli*^49^. These CORE cycles are proposed to be able to reduce CO_2_ (also at ambient concentrations) to formate, which would allow for further coupling for formate assimilation routes, effectively generating CO_2_ fixation routes via formate as intermediate. The experimentally validated CORE variant relies on the promiscuous activity of a β-keto-acid cleavage enzyme (BKACE) and yields one molecule of formate from one molecule of bicarbonate with the input of one NADPH and 1-3 ATP. While the CORE cycle was originally proposed as a photorespiration bypass, the resulting formate could be fully coupled to growth via a formatotrophic pathway such as the serine cycle, which can also operate at ambient CO_2_, unlike the reductive glycine pathway^50^. We have modeled the resulting pathway to compare it with our proposed CO_2_ fixation cycles (Fig. 6). Based on our results, the CORE+Serine cycle (Fig. S9) shows a comparable yield to the CBB in high CO_2_ and a roughly 20% higher yield in ambient CO_2_. Assuming the limiting BKACE activity can be improved, the CORE+Serine cycle could be a highly relevant alternative to conventional CO_2_ fixation cycles, since it is oxygen-tolerant, does not require B_12_, and mostly relies on reactions that are natively present in plants, or that have been shown to be functional after heterologous expression.

## Discussion

In this paper, we proposed several novel carbon fixation pathways based on non-natural enzymatic reactions. We analyzed the pathways based on thermodynamic driving force, predicted biomass yield, and predicted pathway activity. This last parameter is computed based on the measured or estimated k_cat_ and K_M_ values for the enzymes. Unfortunately, the available enzymatic data in databases such as BRENDA is both limited and noisy, since different authors perform enzymatic assays in wildly different conditions in terms of pH, buffers, temperatures, etc. We carefully curated the chosen values, also based on the previous work by Lowe and Kremling^57^, however a relatively large fraction of the enzymes was only measured in single studies, reducing reliability. Where multiple data points are available, it is not unusual to see a variation of one order of magnitude or more in reported k_cat_ and K_M_.

Furthermore, in vitro kinetics measurements, even when performed properly, are based on a purified enzyme in a dilute buffer, which is a completely different scenario compared to the densely packed cytosol of living organisms, where thousands of enzymes and metabolites are present. Recent work has shown k_cat_ measurements in vivo based on different growth conditions and proteomics data^58^. These results show that while many in vitro measurements are also accurate in vivo, there are several outliers even for well characterized enzymes in central metabolism, with a relatively weak correlation R^2^ of 0.62. All these problems are compounded when it comes to relatively understudied enzymes, and especially their promiscuous activities. Finally, the kinetic parameters of not yet existing enzymes are obviously impossible to obtain, and we have simply taken representative values for similar enzymes on related substrates.

In this work, we used the ECM (enzyme cost minimization) method to predict pathway kinetics. This method assumes an optimal distribution of metabolites and enzymes, and does not take into account enzyme degradation, allosteric inhibition, competitive inhibition, and regulation. It also does not consider all the enzymes that are necessary for cell maintenance and growth, assuming all flux is going via the pathway of interest to generate the desired product. In practice, all these factors mean that the predicted pathway activity is at best an upper bound for the real pathway activity in a real-life scenario. Taking the CBB into account, measured values for CO_2_ fixation rates in cell extracts are in the order of 3-9 nmol/min/mg (total) protein^59^. This is a two order of magnitude difference below our computed value of around 0.6 µmol/min/mg (pathway) protein. This discrepancy is partially explained when considering the fact that CBB enzymes account for about 10% of total protein^60^. Additionally, the measured values are from in vitro experiments and likely underestimate the real in vivo value. Based on the Calvin cycle enzyme content of bacterial cells growing at a typical autotrophic doubling time of 4 hours, the expected value for the rate of carbon fixation is 0.25 µmol/min/mg, which is in the same range as the ECM predicted activity for the CBB (see Methods section for the calculations).

For the other pathways, greater uncertainty is expected due to the limited reliability of the available data. It is unclear whether this uncertainty leads to an overestimate or an underestimate of the total pathway activity, however, it is likely that for most of the candidate enzymes, kinetically superior isoforms could be found through a screening process. This points to the fact that pathways other than the CBB may be underestimated in our analysis. Further research will be needed to determine real life (i.e. in vivo) kinetic parameters for several enzymes, leading to better estimates.

We also note that stoichiometric, thermodynamic, and kinetic analyses do not take into account potential issues with some exotic or nonstandard metabolites. These may interact with existing enzymes either as competitive substrates, inhibitors, or regulators, which would require additional engineering efforts (e.g. gene deletions, enzyme engineering) or adaptive laboratory evolution. Importantly, this is not an issue that is specific to the pathways presented in this work, as it commonly arises when engineering synthetic pathways containing metabolites that are not native to the organism of choice^13,61^.

Considering all of this, our proposed pathways are especially promising for potential future applications in terms of yield, as they can outperform the CBB cycle. All the 6 proposed pathways could potentially outperform the yield of the CBB cycle at ambient CO_2_, whereas only the DRED cycle is predicted to outperform the CBB in terms of pathway activity at high CO_2_. Also, the β-glutamate, 3HGL-MC and DRED cycles have relatively high yield gains versus the CBB cycle at ambient CO_2_.

Overall, this makes the DRED cycle promising to explore, given its attractive theoretical yield and predicted pathway activity. However, given the required three novel activities, this cycle would be a tremendous engineering effort already in bacteria, and even more so in microalgae or plants. The ABC-CCL and 3HGL-MC cycles may strike a balance between yield, activity, and relative ease of implementation, requiring only one engineered enzyme.

The RED cycle has a more moderate yield advantage at ambient CO_2_ (15% vs. CBB cycle) but could also be relevant to explore based on its overlap with the CBB cycle and high predicted pathway activity, however it requires the highly challenging ribulose-5-phosphate carboxylase activity, which is shared with the DRED cycle.

The best pathways in this work are predicted to have comparable performance to the best previously proposed mix-and-match pathways, notably the CETCH and THETA cycles^6,54^. However, both these cycles require B_12_-dependent mutases, unlike the cycles proposed in this work. Hence, our proposed cycles have a higher potential for implementation in microalgae and plants, assuming the missing enzymes can be engineered.

The fact that none of the proposed pathways massively outperform the CBB cycle (at high CO_2_) also highlights the fact that carbon fixation cycles are not exclusively limited by carboxylation activity and that Rubisco, despite its relatively slow catalysis and oxygenation side activity, still leads to relatively high pathway activity when considered in its metabolic context^62^.

Still, an open question remains for the field: why is the CBB ubiquitous, if stoichiometrically and/or kinetically superior pathways are potentially possible? One possible explanation for this is the high degree of overlap between the CBB cycle and the pentose phosphate pathway, which exists in all living organisms. In fact, only Rubisco and PRK are necessary on top of the standard pentose phosphate pathway genes, as demonstrated by the engineering of *E. coli* to grow autotrophically via the CBB cycle, and it is common in nature for these two genes to be expressed from a plasmid, which can easily be horizontally transferred, thus spreading the CBB cycle across phyla. Furthermore, several essential biomass precursors are formed as part of the cycle, which would make a loss of activity detrimental. At least in the short term, an organism that acquired all the necessary genes to support an alternative carbon fixation cycle pathway would have to support both cycles at the same time, likely with competing if not outright incompatible fluxes and metabolic profiles.

Even though the proposed pathways likely did not emerge naturally, the well-demonstrated metabolic engineering approach to rewire metabolism using auxotrophies and bypasses to force metabolic rearrangements may allow their realization^63^. This strategy has proven highly reliable in engineering a diverse array of pathways and will likely be just as useful in the future^40,64–67^.

Overall, our study expands the solution space for carbon fixation by providing biochemically plausible avenues for enzymatic transformations based on known enzyme mechanisms and substrate promiscuity. Notably, we identified two potential novel carboxylases that could lead to entirely new pathway architectures if implemented. New carbon fixation pathways with better yield and insensitivity to oxygen could lead to improved carbon fixation in algae or plants, potentially leading to more efficient renewable production of chemicals from CO_2_ and increases in crop yields.

## Acknowledgements

The authors would like to thank Dr. Beau Dronsella for helpful comments on the manuscript. VR is funded by the VLAG Graduate School Wageningen University, NJC acknowledges funding from the ERC StG FASTFIX by the European Research Council and a Vidi grant (VI.Vidi.233.193) from the Dutch Science Organisation (NWO).

## Methods

### Bioretrosynthetic pathway design

Novel carbon fixation pathways were generated through an iterative, expert-guided bioretrosynthetic exploration of biochemical reaction space. Pathway generation was performed manually by considering possible biochemical transformations that could connect metabolites in a carbon fixation cycle. Two complementary starting points were used. In some cases, a desired pathway product or intermediate (e.g. GAP, acetyl-CoA, glyoxylate) was selected and possible biosynthetic precursors were identified recursively. Alternatively, a candidate carbon fixation reaction was selected first, after which downstream and upstream reactions were identified until a closed pathway was obtained.

For each metabolite, candidate transformations were identified based on three levels of biochemical evidence (Table S2). First, reactions that have been experimentally demonstrated for the same substrate or a closely related substrate were considered directly. Second, reactions were proposed by analogy to known enzymatic transformations when the underlying chemistry and catalytic mechanism were sufficiently similar. For example, an aminomutase reaction on a substrate for which no natural activity has been reported may be considered based on the existence of aminomutases catalyzing analogous transformations on other amino acids. Third, transformations were considered when they were chemically plausible based on general principles of enzymatic catalysis, even when no natural example of the specific reaction was known. Examples include the extension of biotin-dependent carboxylation chemistry to alternative enolate substrates and the oxidation or reduction of carbonyl groups on substrates for which a corresponding natural enzyme has not been identified. These latter reactions were considered candidate enzyme activities rather than established biochemical reactions.

The retrosynthetic procedure was applied recursively to each newly identified precursor. As an example, glyoxylate can be connected retrosynthetically to malyl-CoA, which can in turn be derived from malate or fumaryl-CoA. Continuing this procedure can connect malate to oxaloacetate, oxaloacetate to pyruvate, and subsequently through alanine, β-alanine, malonate semialdehyde, malonyl-CoA, and acetyl-CoA, ultimately generating a closed carbon fixation cycle (i.e. the MOG-BAAM cycle). Thus, pathway generation did not require all individual reactions to be naturally occurring, but instead allowed known reactions to be combined with plausible but unreported enzyme activities.

The exploration of reaction space was not intended to be an exhaustive enumeration of all theoretically possible biochemical pathways. Instead, candidate branches were prioritized based on biochemical plausibility and the expected feasibility of the resulting pathway. Branches were generally pursued when the proposed transformation was supported by a known reaction or mechanistic analogy, could plausibly be achieved through enzyme engineering, or represented a chemically reasonable extension of an established catalytic mechanism. Branches were deprioritized when the required reaction appeared unlikely to proceed at a useful rate, when reaching a closed cycle required an impractically large number of additional transformations, or when alternative transformations provided a more favorable route based on considerations such as reaction chemistry, pathway length, or cofactor requirements. Consequently, the resulting pathways represent selected solutions within the broader latent biochemical reaction space rather than an exhaustive set of possible pathways.

For each proposed non-natural reaction, the feasibility of obtaining an appropriate enzyme was considered separately from the feasibility of the overall pathway. Where a closely related natural activity exists, the proposed reaction was considered a potential enzyme-engineering target. For example, carboxylation of α-ketobutyrate was considered plausible based on its close structural similarity to pyruvate, the native substrate of pyruvate carboxylase. In other cases, greater differences between the proposed and known substrates or reactions imply that substantial protein engineering or de novo enzyme design may be required.

The resulting pathways were subsequently evaluated computationally for stoichiometric performance, thermodynamic driving force, enzyme cost, and predicted pathway activity as described below. Pathways were also assessed for requirements for oxygen-sensitive enzymes, vitamin B_12_, and other cofactors or specialized enzyme properties.

### Computational analysis

All scripts and input files are available on gitlab (https://git.wur.nl/vittorio.rainaldi/lvl4), along with all MDF and ECM results. All pathways were computationally analyzed using flux balance analysis^68^ (FBA) with cobrapy^69^ (version 0.29.1) using the *E. coli* iML1515^11^ genome scale-metabolic model. The base model was modified to change membrane-bound transhydrogenase (TDH2pp) to translocate one proton rather than two^70^. Anaerobic reactions such as PFL, OBTFL, and POR5 were set to zero. Non-growth associated maintenance (ATPM) was set to zero. Additional modifications were added for each individual pathway. FBA returns a flux value in gCDW/mol substrate. In this case, since CO_2_ is freely available in the atmosphere, it was left unconstrained, however an energy source is required to run CO_2_ fixation cycles, and this was used as a constraint for all pathways. For this purpose, we added a dummy reaction which produces NADPH, which is the main source of reducing power in photosynthetic organisms such as algae and plants. This results in slightly higher values compared to using NADH as an energy source, since NADH to NADPH conversion requires the translocation of one proton, which amounts to roughly one-third of an ATP^70^.

Max-min driving force^10^ and enzyme cost^12^ were computed with a custom script modified from previous work^44,71^ using equilibrator (version 0.6.0). Metabolite concentrations were constrained to the physiologically relevant range between 10 µM and 10 mM. Ambient CO_2_ is constrained between 10 and 20 µM. To model the CBB cycle, we assumed a 20% rate of Rubisco oxygenation at ambient CO_2_, and no oxygenation at high CO_2_ (3.4 mM, corresponding to a partial pressure of 10%, where photorespiration is virtually absent). ECM was computed using the following parameters: algorithm ECM, version 3, kcat_source fwd, denominator CM, regularization volume, p_h 7.5, ionic_strength 0.3 molar, p_mg 3, dg_confidence 0.95. Kinetics values were taken from Brenda^72^ or previous work^7^. In the case of non-natural enzymatic activities, the kinetics were estimated by taking values from similar enzymes. For example, kinetic values for α-ketobutyrate carboxylase are not available, so values for pyruvate carboxylase were used instead. Since this enzyme catalyzes the same reaction on a slightly smaller substrate, it is reasonable to assume that an engineered enzyme could reach similar kinetic properties. The ECM algorithm returns a list of enzyme costs in moles of enzyme per liter based on a total flux through the pathway for 1 mM/s of product formed. These values were multiplied by the molecular weight of the respective enzyme to obtain the enzyme cost in g/l. The sum of all the values is the enzyme cost (EC) for the pathway. To obtain the pathway activity we performed the following calculations:

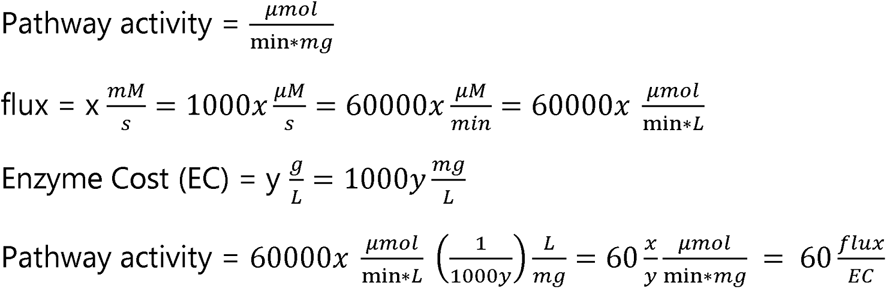

Our ECM results indicate a production rate of around 0.84 µmol/min/mg for the CBB cycle, which is equivalent to carbon fixation rate of 2.5 µmol/min/mg total protein. Assuming pathway enzymes are limited to 10% of the total proteome^73,74^, the rate of carbon fixation becomes 0.25 µmol/min/mg. We can also estimate the rate of carbon fixation needed for growth from a carbon content of 0.5 g/g (Bionumber 100595) and a carbon molecular weight of 12 g/mol. The cell carbon content is thus around 41.6 mmol/gCDW. Based on this number, and a protein content of 0.5 g/g, the carbon content per mg of protein is 41.6 * 0.001 / 0.5 = 0.00832 mmol/mg, or 83.2 µmol/mg. The rate of carbon fixation necessary to support growth at a certain doubling time is given by 83.2 * ln(2)/DT. For a typical autotrophic DT of 4h^75^, this results in a carbon fixation rate of around 14.4 µmol/h/mg, or 0.24 µmol/min/mg, which is close to the ECM prediction

### ECM sensitivity analysis

The sensitivity of enzyme cost minimization (ECM) estimates to uncertainty in kinetic parameters was assessed using Monte Carlo sampling. For each pathway, k_cat_ and K_M_ specified in the input parameter table were independently perturbed according to a log-normal multiplicative model.

For each pathway, the unperturbed model was solved first to obtain the baseline cost. Then, N = 1000 perturbed parameter sets were drawn. Each k_cat_ and K_M_ value was multiplied by a random factor drawn from a log-normal distribution with median 1, i.e. ln(m) ∼ Normal(0, σ²). The spread σ was set directly as a typical fold change, σ = ln(f). A value of f = 2 was used, so 68% of parameters (1 SD) are between half and double the original value, and 95% of draws fall within ∼¼ and ∼4 times. One multiplier was drawn per reaction and applied to all k_cat_ entries of that reaction (forward, reverse and geometric mean). An independent multiplier was drawn for every K_M_ entry. Standard Gibbs energies, flux values, concentration bounds, and molecular masses were held fixed, and parameters were sampled independently of each other.

For each pathway, the total enzyme cost calculated using the unperturbed parameter set was used as a reference. The distribution of costs obtained from successful Monte Carlo samples was summarized using the mean, median, 5th and 95th percentiles, and coefficient of variation. The unperturbed cost and Monte Carlo median and mean were also indicated in histograms of the sampled total enzyme costs. For comparisons among pathways, results were ranked by the mean total enzyme cost. The script and results are available in the online repository associated with this study. Results are also summarized in Table S3.

## Notes

https://git.wur.nl/vittorio.rainaldi/lvl4

### Competing Interest Statement

The authors have declared no competing interest.

### Summary of Updates

Added updates to kinetics analysis and additional tables. Revised main text.

## References

1. The Prokaryotes. (Springer Berlin Heidelberg, Berlin, Heidelberg, 2013). doi:10.1007/978-3-642-30123-0.

2. Prywes, N., Phillips, N. R., Tuck, O. T., Valentin-Alvarado, L. E. & Savage, D. F. Rubisco Function, Evolution, and Engineering. Annu. Rev. Biochem. 92, 385–410 (2023).

3. Tcherkez, G. G. B., Farquhar, G. D. & Andrews, T. J. Despite slow catalysis and confused substrate specificity, all ribulose bisphosphate carboxylases may be nearly perfectly optimized. Proceedings of the National Academy of Sciences 103, 7246–7251 (2006).

4. Claassens, N. J., et al. Phosphoglycolate salvage in a chemolithoautotroph using the Calvin cycle. Proc. Natl. Acad. Sci. U.S.A. 117, 22452–22461 (2020).

5. Bar-Even, A., Noor, E., Lewis, N. E. & Milo, R. Design and analysis of synthetic carbon fixation pathways. Proceedings of the National Academy of Sciences 107, 8889–8894 (2010).

6. Schwander, T., Von Borzyskowski, L. S., Burgener, S., Cortina, N. S. & Erb, T. J. A synthetic pathway for the fixation of carbon dioxide in vitro. Science 354, 900–904 (2016).

7. Löwe, H. & Kremling, A. In-Depth Computational Analysis of Natural and Artificial Carbon Fixation Pathways. BioDesign Research 2021, 1–23 (2021).

8. Vornholt, T. et al. Artificial metalloenzymes. Nat Rev Methods Primers 4, 78 (2024).

9. Zou, Z. et al. De novo design and evolution of an artificial metathase for cytoplasmic olefin metathesis. Nat Catal 1–12 (2025) doi:10.1038/s41929-025-01436-0.

10. Noor, E. et al. Pathway Thermodynamics Highlights Kinetic Obstacles in Central Metabolism. PLoS Computational Biology 10, (2014).

11. Monk, J. M. et al. iML1515, a knowledgebase that computes Escherichia coli traits. Nat Biotechnol 35, 904–908 (2017).

12. Noor, E. et al. The Protein Cost of Metabolic Fluxes: Prediction from Enzymatic Rate Laws and Cost Minimization. PLoS Computational Biology 12, 1–29 (2016).

13. Rainaldi, V. et al. Modular *in vivo* engineering of the reductive methylaspartate cycles for synthetic CO2 fixation. Trends in Biotechnology 10.1016/j.tibtech.2026.09.019 (2026) doi:10.1016/j.tibtech.2026.09.019.

14. Croft, M. T., Lawrence, A. D., Raux-Deery, E., Warren, M. J. & Smith, A. G. Algae acquire vitamin B12 through a symbiotic relationship with bacteria. Nature 438, 90–93 (2005).

15. Zhu, X. G., Long, S. P. & Ort, D. R. What is the maximum efficiency with which photosynthesis can convert solar energy into biomass? Current Opinion in Biotechnology 19, 153–159 (2008).

16. Walker, B. J., VanLoocke, A., Bernacchi, C. J. & Ort, D. R. The Costs of Photorespiration to Food Production Now and in the Future. Annu. Rev. Plant Biol. 67, 107–129 (2016).

17. Sharkey, T. D. Estimating the rate of photorespiration in leaves. Physiologia Plantarum 73, 147–152 (1988).

18. Alban, C., Baldet, P., Axiotis, S. & Douce, R. Purification and Characterization of 3-Methylcrotonyl-Coenzyme A Carboxylase from Higher Plant Mitochondria. Plant Physiol. 102, 957–965 (1993).

19. Huang, C. S., Ge, P., Zhou, Z. H. & Tong, L. An unanticipated architecture of the 750-kDa α6β6 holoenzyme of 3-methylcrotonyl-CoA carboxylase. Nature 481, 219–223 (2012).

20. Chavez-Avilas, M., Díaz-Pérez, A. L., Reyes-de la Cruz, H. & Campos-García, J. The *Pseudomonas aeruginosa liuE* gene encodes the 3-hydroxy-3-methylglutaryl coenzyme A lyase, involved in leucine and acyclic terpene catabolism. FEMS Microbiology Letters 296, 117–123 (2009).

21. Wang, H., et al. An essential bifunctional enzyme in *Mycobacterium tuberculosis* for itaconate dissimilation and leucine catabolism. Proc. Natl. Acad. Sci. U.S.A. 116, 15907–15913 (2019).

22. Chu, C.-H. & Cheng, D. Expression, purification, characterization of human 3-methylcrotonyl-CoA carboxylase (MCCC). Protein Expression and Purification 53, 421–427 (2007).

23. Raj, H. et al. Engineering methylaspartate ammonia lyase for the asymmetric synthesis of unnatural amino acids. Nature Chem 4, 478–484 (2012).

24. Herrmann, G. et al. Two beta-alanyl-CoA:ammonia lyases in Clostridium propionicum. The FEBS Journal 272, 813–821 (2005).

25. Ruzicka, F. J. & Frey, P. A. Glutamate 2,3-aminomutase: A new member of the radical SAM superfamily of enzymes. Biochimica et Biophysica Acta (BBA) - Proteins and Proteomics 1774, 286–296 (2007).

26. Tunçkanat, T. et al. Lysine 2,3-Aminomutase and a Newly Discovered Glutamate 2,3-Aminomutase Produce β-Amino Acids Involved in Salt Tolerance in Methanogenic Archaea. Biochemistry 61, 1077–1090 (2022).

27. Chen, D., Ruzicka, F. J. & Frey, P. A. A novel lysine 2,3-aminomutase encoded by the yodO gene of Bacillus subtilis : characterization and the observation of organic radical intermediates. 11 (2000).

28. Chen, D., Frey, P. A., Lepore, B. W., Ringe, D. & Ruzicka, F. J. Identification of Structural and Catalytic Classes of Highly Conserved Amino Acid Residues in Lysine 2,3-Aminomutase. Biochemistry 45, 12647–12653 (2006).

29. Behshad, E. et al. Enantiomeric Free Radicals and Enzymatic Control of Stereochemistry in a Radical Mechanism: The Case of Lysine 2,3-Aminomutases. Biochemistry 45, 12639–12646 (2006).

30. Sofia, H. J., Chen, G., Hetzler, B. G., Reyes-Spindola, J. F. & Miller, N. E. Radical SAM, a novel protein superfamily linking unresolved steps in familiar biosynthetic pathways with radical mechanisms: functional characterization using new analysis and information visualization methods. Nucleic Acids Res 29, 1097–1106 (2001).

31. Booker, S. J. & Lloyd, C. T. Twenty Years of Radical SAM! The Genesis of the Superfamily. ACS Bio Med Chem Au 2, 538–547 (2022).

32. Mack, M. et al. 3-Methylglutaconyl-CoA hydratase from Acinetobacter sp. Arch Microbiol 185, 297–306 (2006).

33. Textor, S. et al. Propionate oxidation in Escherichia coli: Evidence for operation of a methylcitrate cycle in bacteria. Archives of Microbiology 168, 428–436 (1997).

34. London, R. E., Allen, D. L., Gabel, S. A. & DeRose, E. F. Carbon-13 Nuclear Magnetic Resonance Study of Metabolism of Propionate by Escherichia coli. Journal of Bacteriology 10.1128/jb.181.11.3562-3570.1999 (1999) doi:10.1128/jb.181.11.3562-3570.1999.

35. Liu, Y. et al. Systematic design and evaluation of artificial CO2 assimilation pathways. Synthetic and Systems Biotechnology 10, 1107–1118 (2025).

36. Kim, J., Hetzel, M., Boiangiu, C. D. & Buckel, W. Dehydration of (*R*)-2-hydroxyacyl-CoA to enoyl-CoA in the fermentation of α-amino acids by anaerobic bacteria. FEMS Microbiol Rev 28, 455–468 (2004).

37. Kim, J., Darley, D. J., Buckel, W. & Pierik, A. J. An allylic ketyl radical intermediate in clostridial amino-acid fermentation. Nature 452, 239–242 (2008).

38. Bahnson, B. J., Anderson, V. E. & Petsko, G. A. Structural Mechanism of Enoyl-CoA Hydratase: Three Atoms from a Single Water Are Added in either an E1cb Stepwise or Concerted Fashion,. Biochemistry 41, 2621–2629 (2002).

39. Hordijk, W. Autocatalytic confusion clarified. Journal of Theoretical Biology 435, 22–28 (2017).

40. Wenk, S. et al. Evolution-assisted engineering of *E. coli* enables growth on formic acid at ambient CO2 via the Serine Threonine Cycle. Metabolic Engineering 88, 14–24 (2025).

41. Scrutton, M. C., Griminger, P. & Wallace, J. C. Pyruvate Carboxylase. Journal of Biological Chemistry 247, 3305–3313 (1972).

42. Scheffen, M. et al. A new-to-nature carboxylation module to improve natural and synthetic CO2 fixation. Nature Catalysis 4, 105–115 (2021).

43. Rainaldi, V., Donati, S., D’Adamo, S. & Claassens, N. J. A recursive pathway for isoleucine biosynthesis arises from enzyme promiscuity. 2025.08.26.672309 Preprint at 10.1101/2025.08.26.672309 (2025).

44. Satanowski, A. et al. Awakening a latent carbon fixation cycle in Escherichia coli. Nature Communications 11, 1–14 (2020).

45. Aoshima, M. & Igarashi, Y. A novel oxalosuccinate-forming enzyme involved in the reductive carboxylation of 2-oxoglutarate in Hydrogenobacter thermophilus TK-6. Molecular Microbiology 62, 748–759 (2006).

46. Aoshima, M. & Igarashi, Y. Nondecarboxylating and Decarboxylating Isocitrate Dehydrogenases: Oxalosuccinate Reductase as an Ancestral Form of Isocitrate Dehydrogenase. J Bacteriol 190, 2050–2055 (2008).

47. Buhrman, G. et al. Structure, Function, and Thermal Adaptation of the Biotin Carboxylase Domain Dimer from Hydrogenobacter thermophilus 2-Oxoglutarate Carboxylase. Biochemistry 60, 324–345 (2021).

48. Bar-Even, A. Daring metabolic designs for enhanced plant carbon fixation. Plant Science 273, 71–83 (2018).

49. Satanowski, A. et al. Design and implementation of aerobic and ambient CO2-reduction as an entry-point for enhanced carbon fixation. Nat Commun 16, 3134 (2025).

50. Cowan, A. E. et al. Fast growth and high-titer bioproduction from renewable formate via metal-dependent formate dehydrogenase in Escherichia coli. Nat Commun 16, 5908 (2025).

51. Yang, X., Ma, Y., Yan, J., Zhao, G. & Zhang, Y. LATCH: a Linear Autocatalytic cycle Tailored for Carbon Harvesting. Green Carbon 10.1016/j.greenca.2025.10.003 (2025) doi:10.1016/j.greenca.2025.10.003.

52. Bar-Even, A., Noor, E., Lewis, N. E. & Milo, R. Design and analysis of synthetic carbon fixation pathways. Proceedings of the National Academy of Sciences of the United States of America 107, 8889–8894 (2010).

53. Schulz-Mirbach, H. et al. New-to-nature CO2-dependent acetyl-CoA assimilation enabled by an engineered B12-dependent acyl-CoA mutase. Nat Commun 15, 10235 (2024).

54. Luo, S. et al. Construction and modular implementation of the THETA cycle for synthetic CO2 fixation. Nat Catal 6, 1228–1240 (2023).

55. McLean, R. et al. Exploring alternative pathways for the in vitro establishment of the HOPAC cycle for synthetic CO _2_ fixation. Sci. Adv. 9, eadh4299 (2023).

56. Luo, S., et al. A cell-free self-replenishing CO2-fixing system. Nat Catal 5, (2022).

57. Löwe, H. & Kremling, A. In-Depth Computational Analysis of Natural and Artificial Carbon Fixation Pathways. BioDesign Research 2021, 9898316 (2021).

58. Davidi, D. et al. Global characterization of in vivo enzyme catalytic rates and their correspondence to in vitro kcat measurements. Proceedings of the National Academy of Sciences of the United States of America 113, 3401–3406 (2016).

59. Gale, N. & Beck, J. Evidence for the Calvin Cycle and Hexose Monophosphate Pathway in Thiobacillus ferrooxidans. https://journals.asm.org/doi/epdf/10.1128/jb.94.4.1052-1059.1967 (1967) doi:10.1128/jb.94.4.1052-1059.1967.

60. Dronsella, B., et al. Engineered Synthetic One-Carbon Fixation Exceeds Yield of the Calvin Cycle. http://biorxiv.org/lookup/doi/10.1101/2022.10.19.512895(2022) doi:10.1101/2022.10.19.512895.

61. Schulz-Mirbach, H. et al. Combining evolution and machine learning-guided pathway optimization to engineer a novel methylsuccinate module for synthetic C1 metabolism in vivo. 2026.01.16.699985 Preprint at 10.64898/2026.01.16.699985 (2026).

62. Bathellier, C., Tcherkez, G., Lorimer, G. H. & Farquhar, G. D. Rubisco is not really so bad. Plant Cell Environ 41, 705–716 (2018).

63. Schulz-Mirbach, H., Dronsella, B. & Erb, T. J. Escherichia coli selection strains for growth-coupled metabolic engineering. Trends in Biotechnology 0, (2025).

64. Gleizer, S. et al. Conversion of Escherichia coli to Generate All Biomass Carbon from CO2. Cell 179, 1255–1263.e12 (2019).

65. Kim, S. et al. Growth of E. coli on formate and methanol via the reductive glycine pathway. Nature Chemical Biology 16, 538–545 (2020).

66. Reiter, M. A. et al. A synthetic methylotrophic Escherichia coli as a chassis for bioproduction from methanol. Nat Catal 7, 560–573 (2024).

67. Chen, F. Y. H., Jung, H. W., Tsuei, C. Y. & Liao, J. C. Converting Escherichia coli to a Synthetic Methylotroph Growing Solely on Methanol. Cell 182, 933–946.e14 (2020).

68. Edwards, J. S., Ibarra, R. U. & Palsson, B. O. In silico predictions of Escherichia coli metabolic capabilities are consistent with experimental data. Nat Biotechnol 19, 125–130 (2001).

69. Ebrahim, A., Lerman, J. A., Palsson, B. O. & Hyduke, D. R. COBRApy: COnstraints-Based Reconstruction and Analysis for Python. BMC Systems Biology 7, 74 (2013).

70. Bizouarn, T., Van Boxel, G. I., Bhakta, T. & Jackson, J. B. Nucleotide binding affinities of the intact proton-translocating transhydrogenase from Escherichia coli. Biochimica et Biophysica Acta - Bioenergetics 1708, 404–410 (2005).

71. Elad Noor / ged-cycle · GitLab. GitLab https://gitlab.com/elad.noor/ged-cycle (2024).

72. Chang, A. et al. BRENDA, the ELIXIR core data resource in 2021: new developments and updates. Nucleic Acids Res 49, D498–D508 (2021).

73. Raven, J. A. Rubisco: still the most abundant protein of Earth? New Phytologist 198, 1–3 (2013).

74. Dronsella, B. et al. One-carbon fixation via the synthetic reductive glycine pathway exceeds yield of the Calvin cycle. Nat Microbiol 1–8 (2025) doi:10.1038/s41564-025-01941-9.

75. Claassens, N. J., Cotton, C. A. R., Kopljar, D. & Bar-Even, A. Making quantitative sense of electromicrobial production. Nature Catalysis 2, 437–447 (2019).

