## Supplementary material for "Design and analysis of synthetic carbon fixation pathways based on novel enzymatic reactions": SI

**Supplementary information**

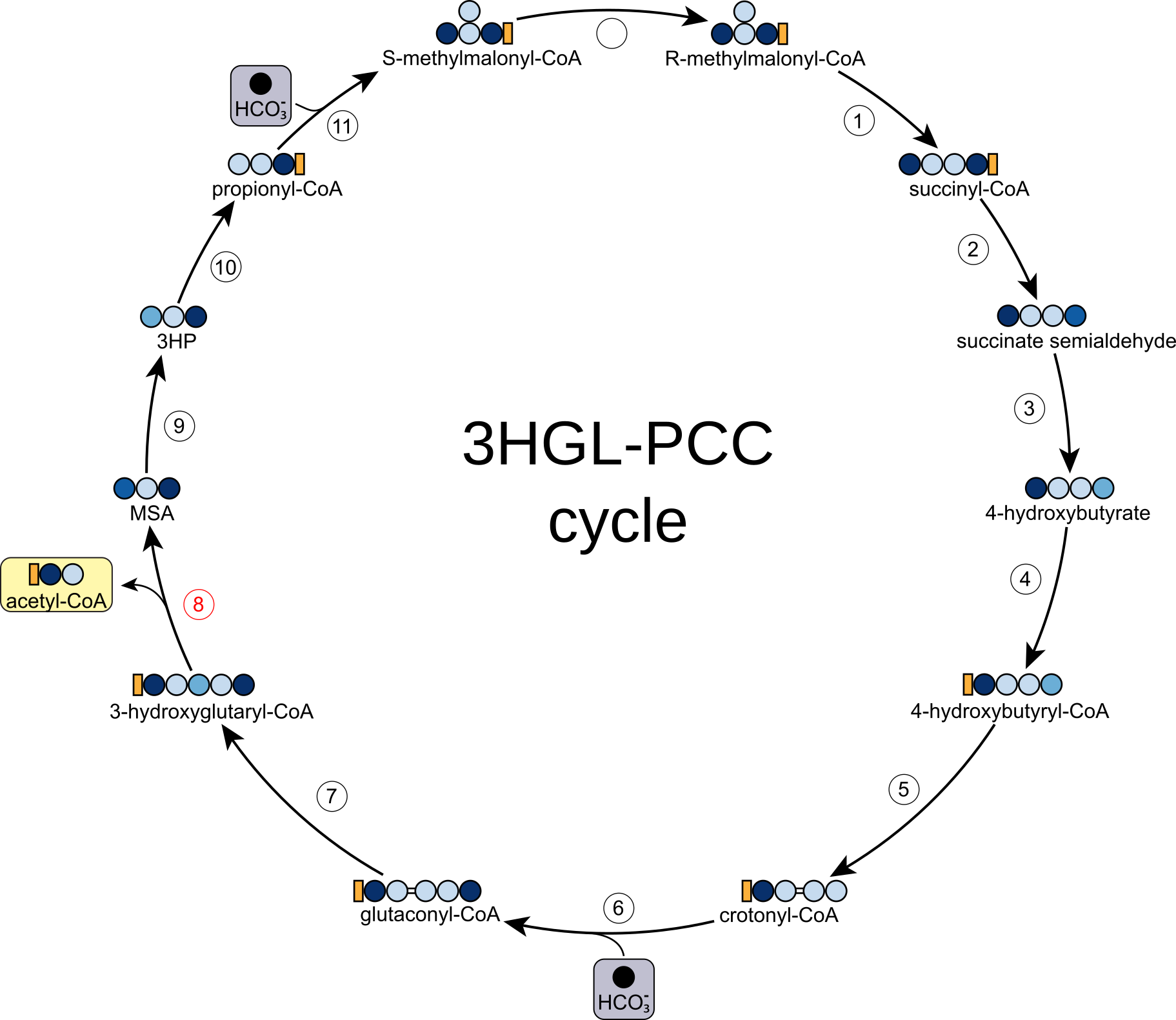

**Figure S1**: The 3-hydroxyglutaryl-CoA lyase-propionyl-CoA carboxylase (3HGL-PCC) cycle. This cycle is a variant of the 3HGL-MC cycle that uses propionyl-CoA carboxylase instead of the methylcitrate cycle combined with pyruvate carboxylase. This results in the production of succinyl-CoA, as opposed to succinate, saving one ATP per cycle and increasing biomass yield. However, the conversion of methylmalonyl-CoA to succinyl-CoA via methylmalonyl-CoA mutase requires B_12_ as a cofactor, limiting the applicability of this cycle. 1, methylmalonyl-CoA mutase; 2-6 are as in figure 1; 7, glutaconyl-CoA hydratase; 8, 3-hydroxyglutaryl-CoA lyase; 9, malonate semialdehyde reductase; 10, propionyl-CoA synthase; 11, propionyl-CoA carboxylase; 12, methylmalonyl-CoA epimerase; 3HP, 3-hydroxypropionate; CoA, coenzyme A; MSA, malonate semialdehyde.

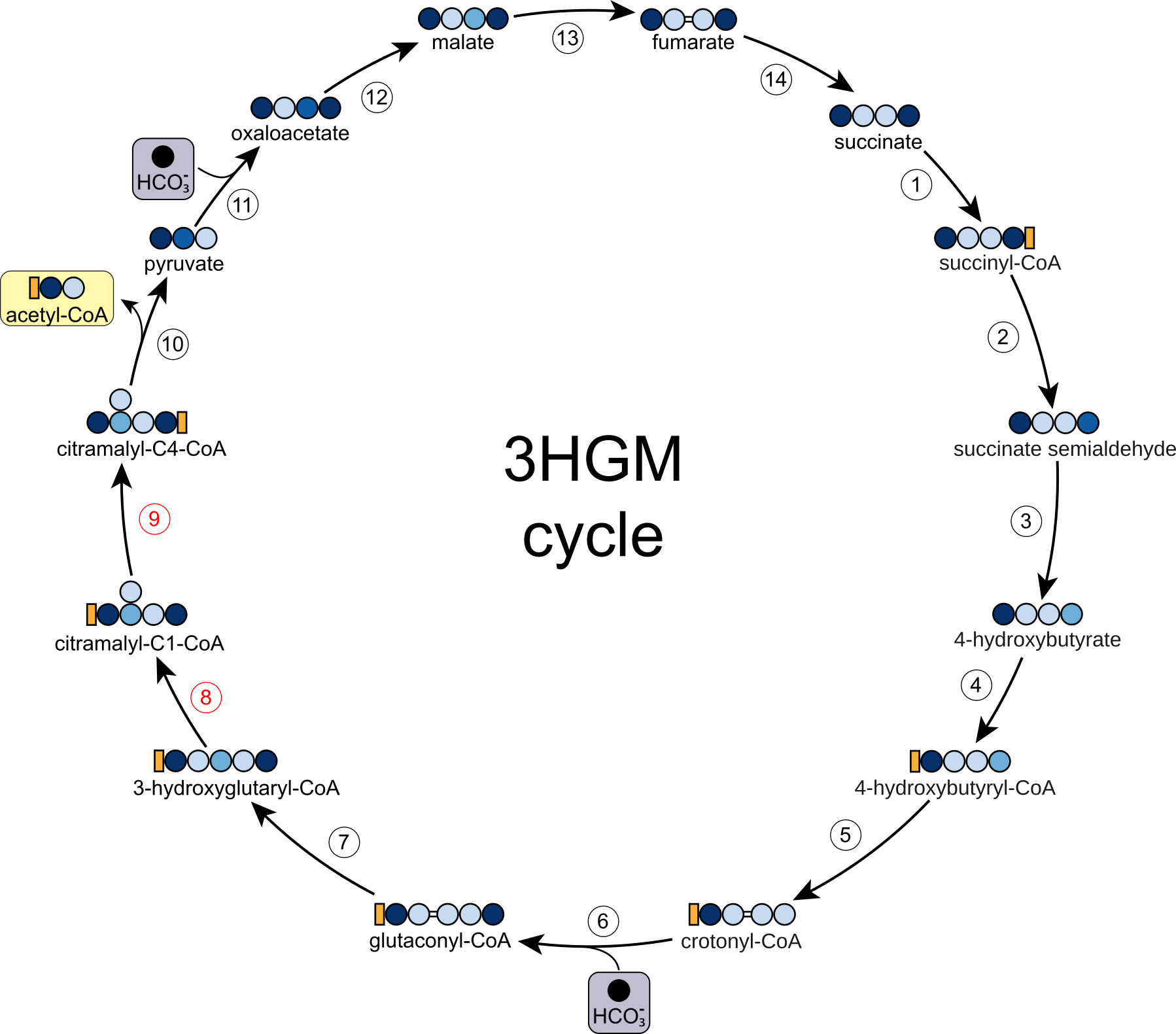

**Figure S2**: The 3-hydroxyglutaryl-CoA mutase cycle. 3-hydroxyglutaryl-CoA formed from the hydration of glutaconyl-CoA could be isomerized in a similar reaction to the naturally occurring interconversion of 3-hydroxybutyryl-CoA and 2-hydroxyisobutyryl-CoA^1,2^, catalyzed by a B_12_-dependent mutase. This would result in the 3HGM cycle, which also has high predicted performance in terms of yield (~30% higher yield at ambient CO_2_ (Table S1). However, it would require two engineered enzymes for which it is unclear if fast enough variants can be realized. The molecule resulting from 3-hydroglutaryl-CoA mutase activity, namely citramalyl-CoA, would carry the CoA on C1 rather than C4, thus requiring a second specialized enzyme to swap the CoA, similarly to mesaconyl-CoA transferase^3^. This is necessary because the subsequent citramalyl-CoA lyase reaction is only possible if the CoA is in C4. It is unclear how fast the 3-hydroxyglutaryl-CoA mutase rate could be. A computational analysis of a similar reaction, namely the promiscuous activity of B_12_-dependent glutamate mutase on the alternative substrate 2-hydroxyglutarate (yielding threo-methylmalate), identified a thermodynamic bottleneck in the reaction due to the formation of a glycolyl radical instead of the usual glycyl radical which is formed in the glutamate mutase reaction. The presence of a hydroxyl group in the glycolyl radical is thought to destabilize the radical leading to an increase in activation energy and a proportional decrease in k_cat_ of about two orders of magnitude (5 s^-1^ to 0.05 s^-1^)^4^. It is currently unclear whether this issue can be solved and if it is related to glutamate mutase or general to all B_12_-dependent mutases. More research will be needed to improve our understanding of the chemistry catalyzed by this complex cofactor. Regardless, this cycle, much like the 3HGL-PCC cycle, is limited by its B_12_ requirement. 1-6 are as in figure 1; 7, glutaconyl-CoA hydratase; 8, 3-hydroxyglutaryl-CoA mutase; 9, citramalyl-CoA transferase; 10, citramalyl-CoA lyase; 11, pyruvate carboxylase; 12, malate dehydrogenase; 13, fumarase; 14, fumarate reductase; CoA, coenzyme A.

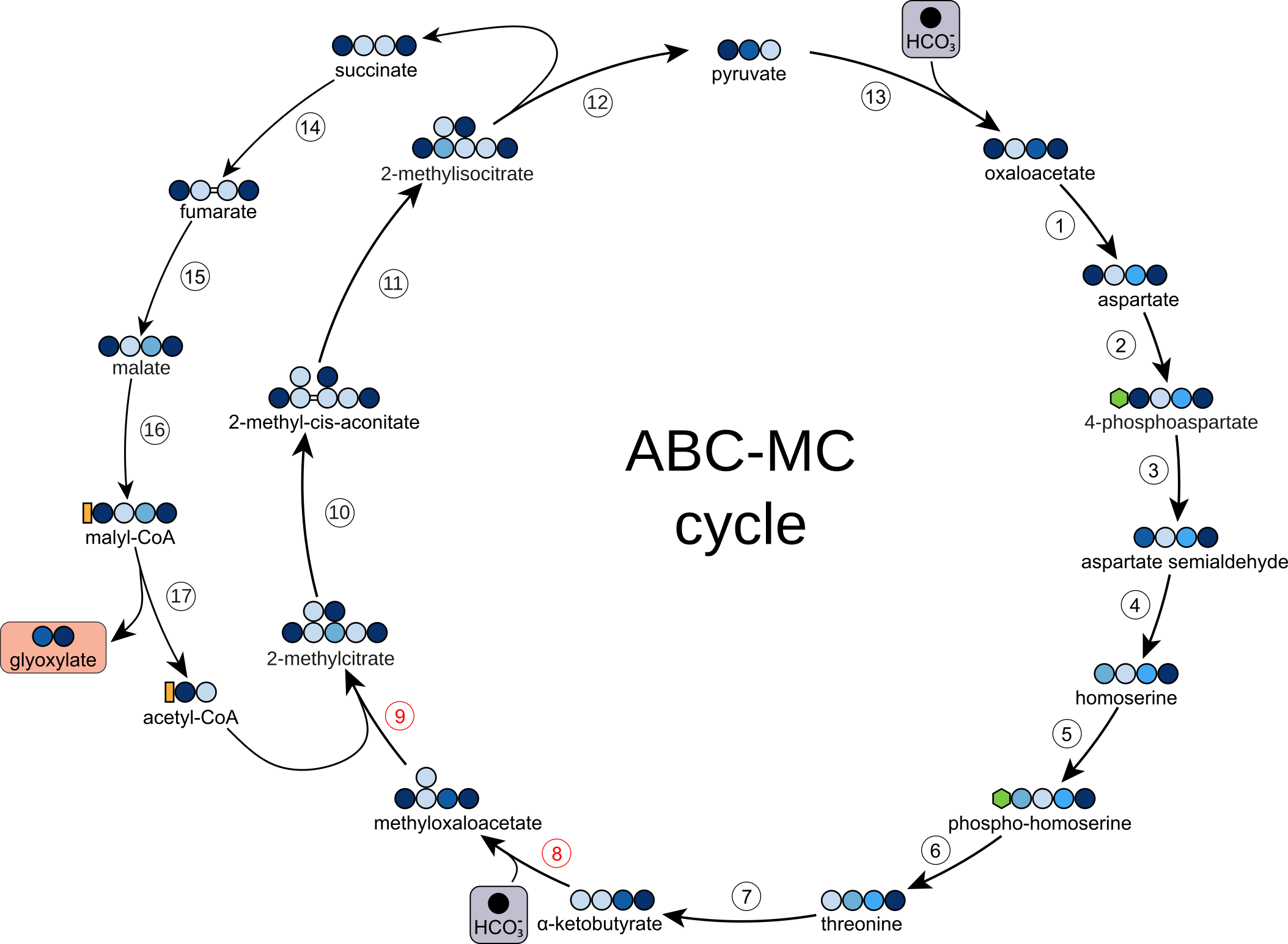

**Figure S3**: The α-ketobutyrate carboxylase-methylcitrate (ABC-MC) cycle. 1-8 are as in figures 3 and 4. 9, methylcitrate synthase; 10, 2-methylcitrate dehydratase; 11, aconitase; 12, methylisocitrate lyase; 13, pyruvate carboxylase; 14, succinate dehydrogenase; 15, fumarase; 16, malate thiokinase; 17, malyl-CoA lyase.

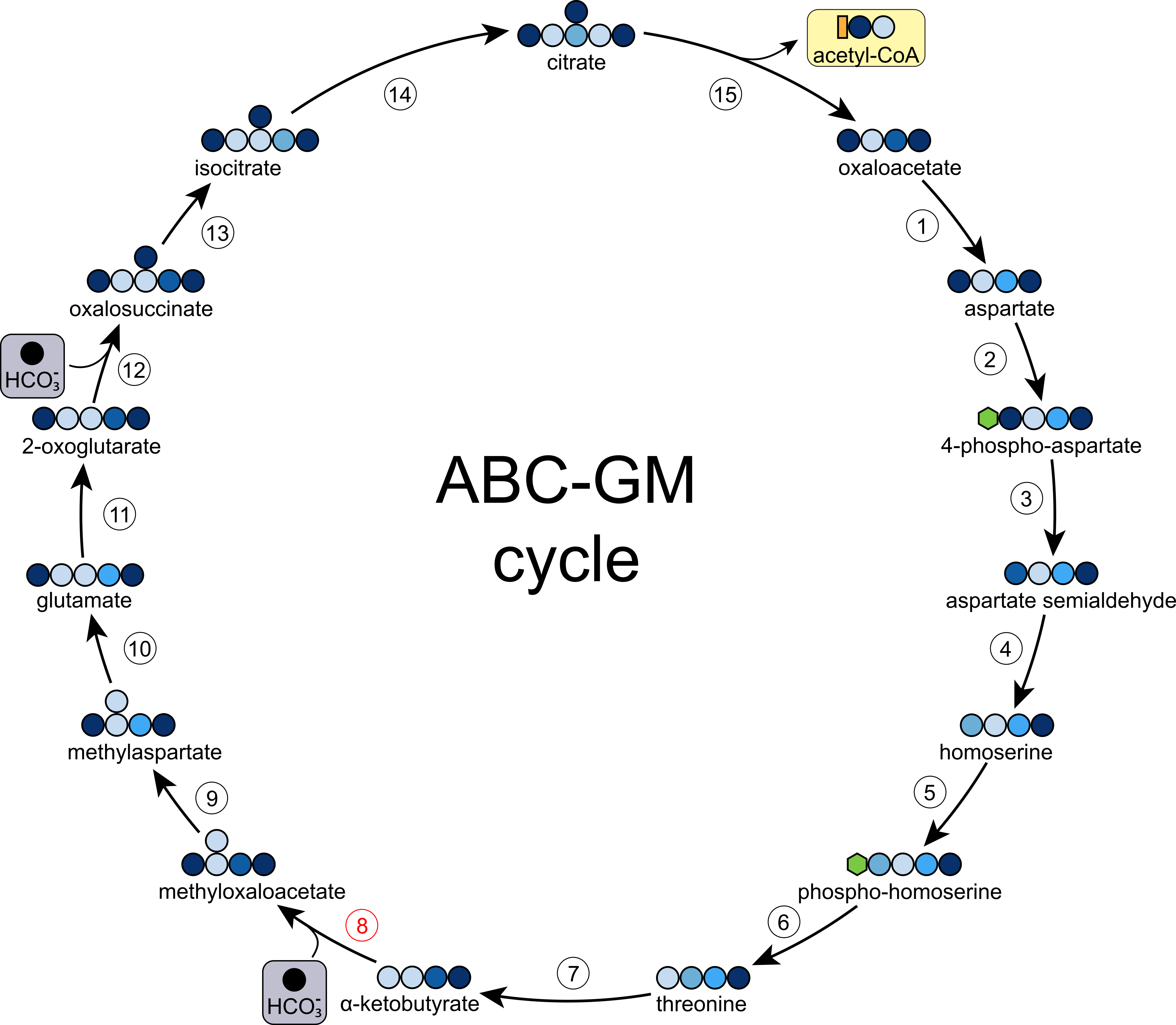

**Figure S4**: The α-ketobutyrate carboxylase-glutamate mutase cycle. 1, aspartate aminotransferase; 2, aspartate kinase; 3, aspartate semialdehyde dehydrogenase; 4, homoserine dehydrogenase; 5, homoserine kinase; 6, threonine synthase; 7, threonine deaminase; 8, *α*-ketobutyrate carboxylase; 9, methylaspartate aminotransferase; 10, glutamate mutase; 11, glutamate dehydrogenase; 12, 2-oxoglutarate carboxylase; 13, oxalosuccinate reductase; 14, aconitase; 15, ATP citrate lyase

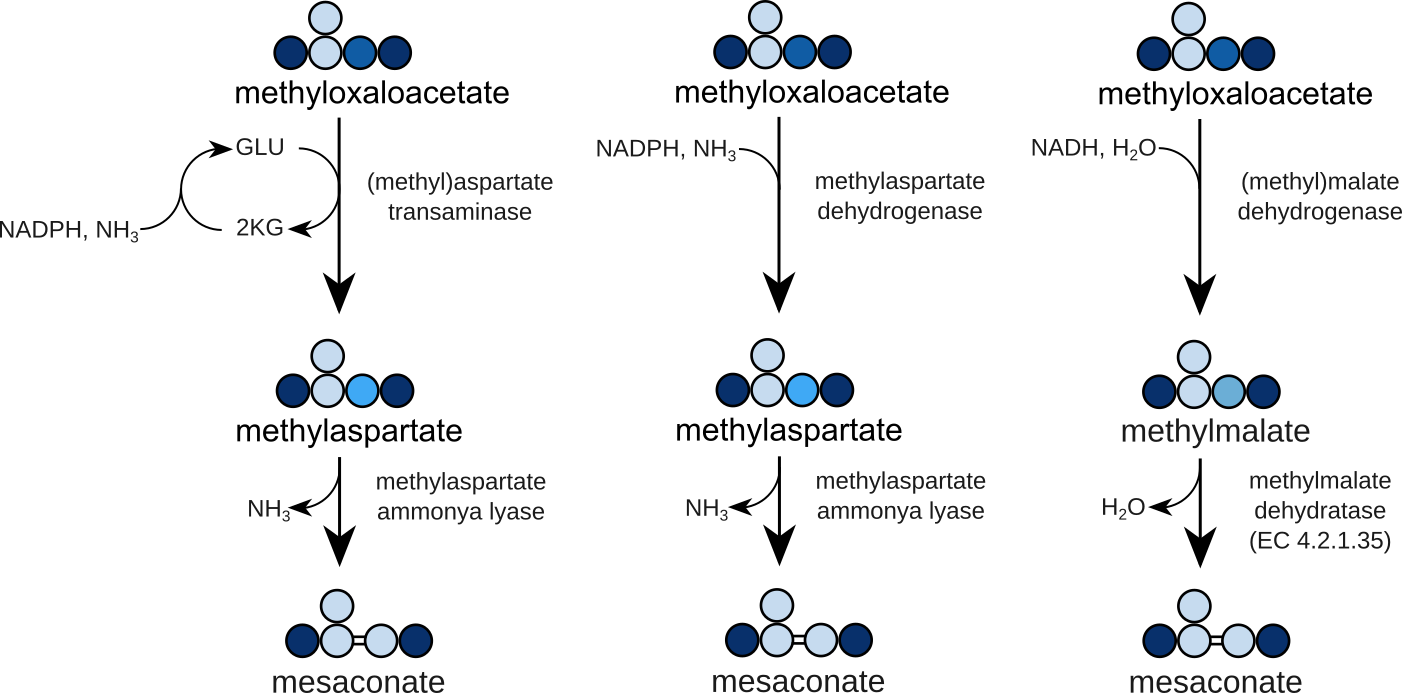

**Figure S5:** Alternative ways to convert methyloxaloacetate to mesaconate. The transaminase route relies on natural enzyme activities, while methylaspartate dehydrogenase and methylmalate dehydrogenase require engineering. Furthermore, while the methylmalate dehydratase reaction is reported on KEGG, we were unable to find a clear source for it, and it could produce citraconate rather than mesaconate, depending on its stereochemistry. The methylmalate dehydrogenase route is expected to be slightly more energy efficient since it uses NADH as a source of reducing power as opposed to NADPH for the other two routes. 2KG, 2-ketoglutarate; GLU, glutamate.

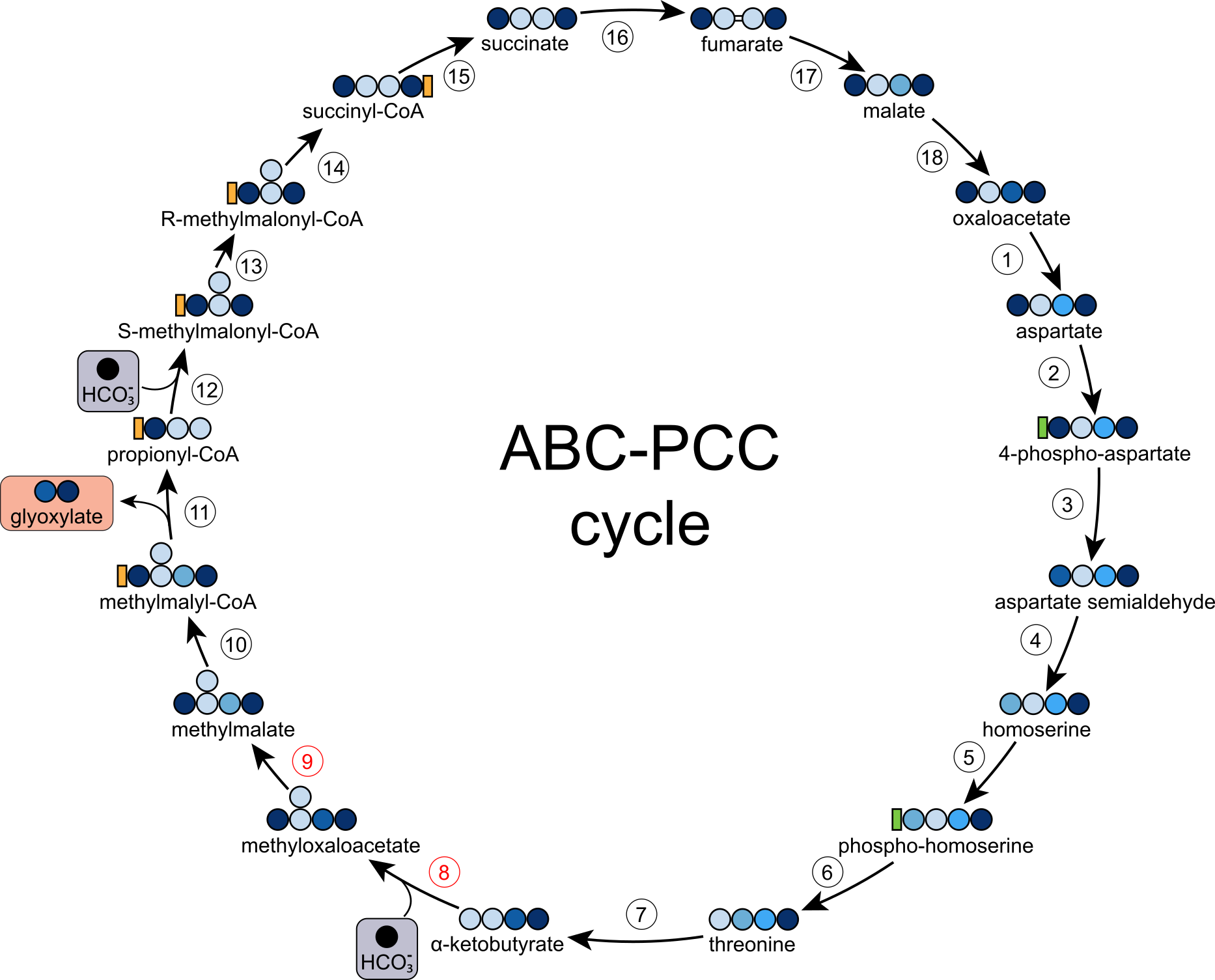

**Figure S6**: The α-ketobutyrate carboxylase-propionyl-CoA carboxylase cycle. 1-8 are as in figures 4 and 5. 9, methylmalate dehydrogenase, 10, methylmalate-CoA transferase; 11, methylmalyl-CoA lyase; 12, propionyl-CoA carboxylase; 13, methylmalonyl-CoA epimerase; 14, methylmalonyl-CoA mutase; 15, succinyl-CoA synthetase; 16, succinate dehydrogenase; 17, fumarase; 18, malate dehydrogenase; CoA, coenzyme A.

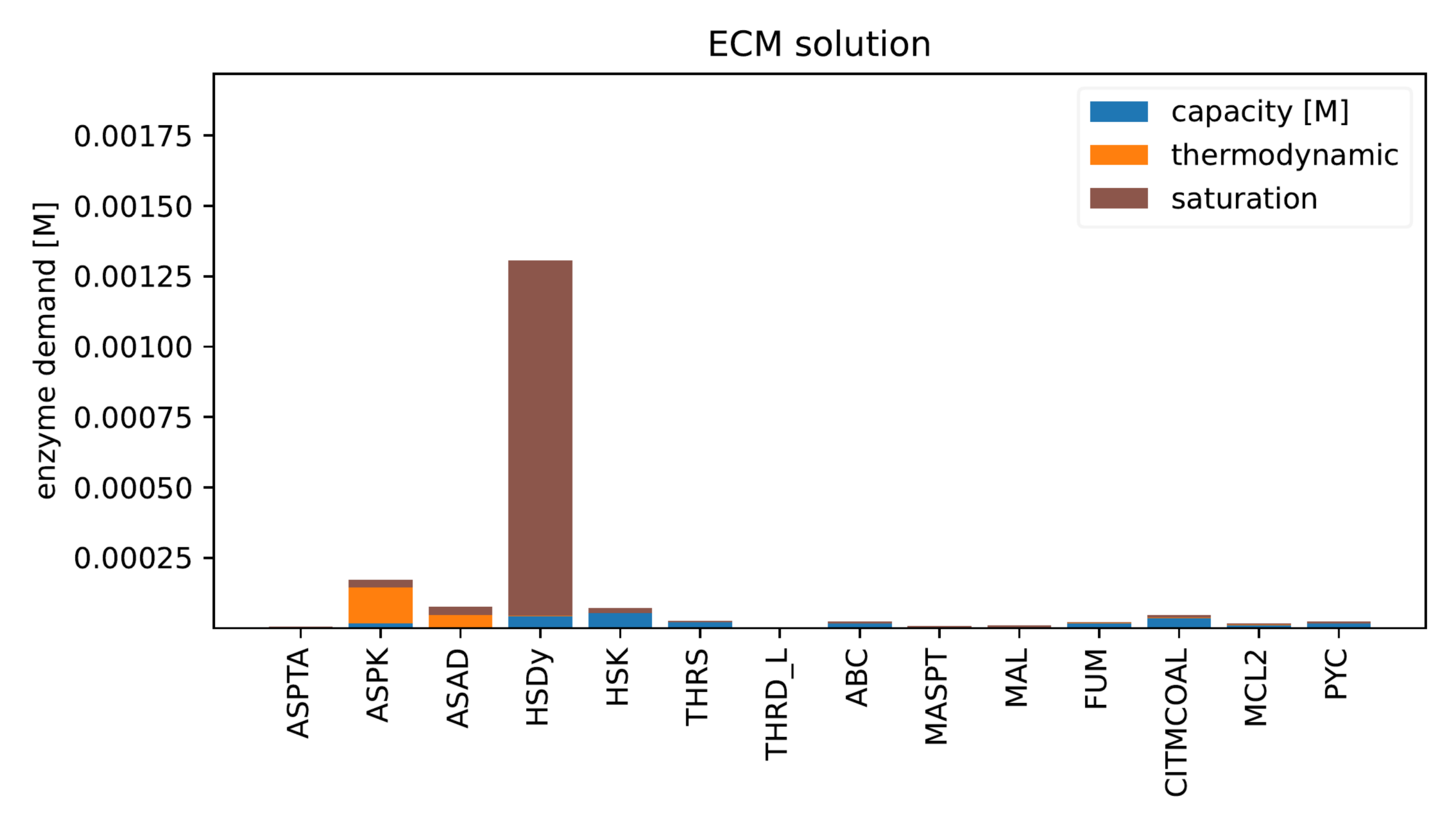
**Figure S7**: ECM costs for each enzyme in the ABC-CCL cycle. According to this computational analysis, homoserine dehydrogenase activity may limit the overall pathway activity. This limitation is due to the inability of the enzyme to be saturated by its substrate under physiological conditions. Since this is a natural enzyme, it may not be possible to improve its kinetic parameters and eliminate the bottleneck. On the other hand, the enzyme may not be optimized to carry a high flux, since it is normally only involved in threonine biosynthesis, which only accounts for roughly 5% of biomass^5^.

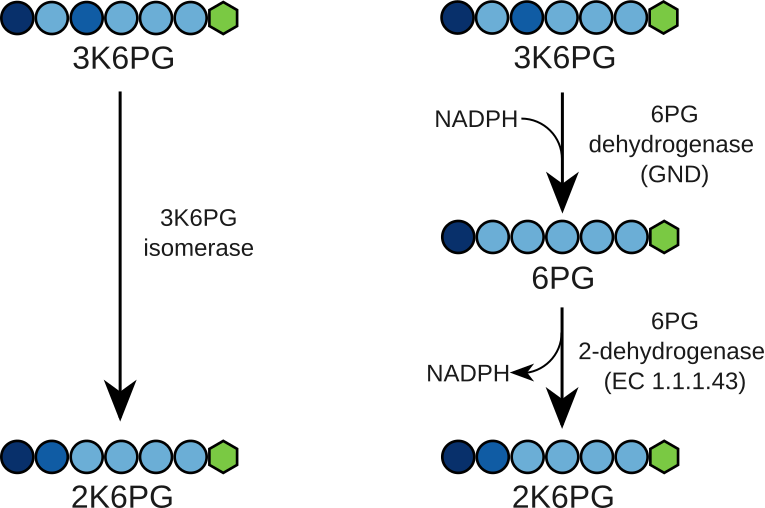

**Figure S8:** Alternative ways to convert 3-keto-6-phosphogluconate to 2-keto-6-phosphogluconate. While the isomerase is not known to exist in nature, both dehydrogenases are known to exist and could provide an easier alternative. Additionally, 3K6PG is a β-ketoacid and this class of compounds tends to spontaneously decarboxylate at neutral pH and ambient temperature. Quickly reducing it to 6PG would prevent this issue, while the product of the following reaction, 2K6PG, is an α-ketoacid that is expected to be fully stable until temperatures well above the boiling point of water.

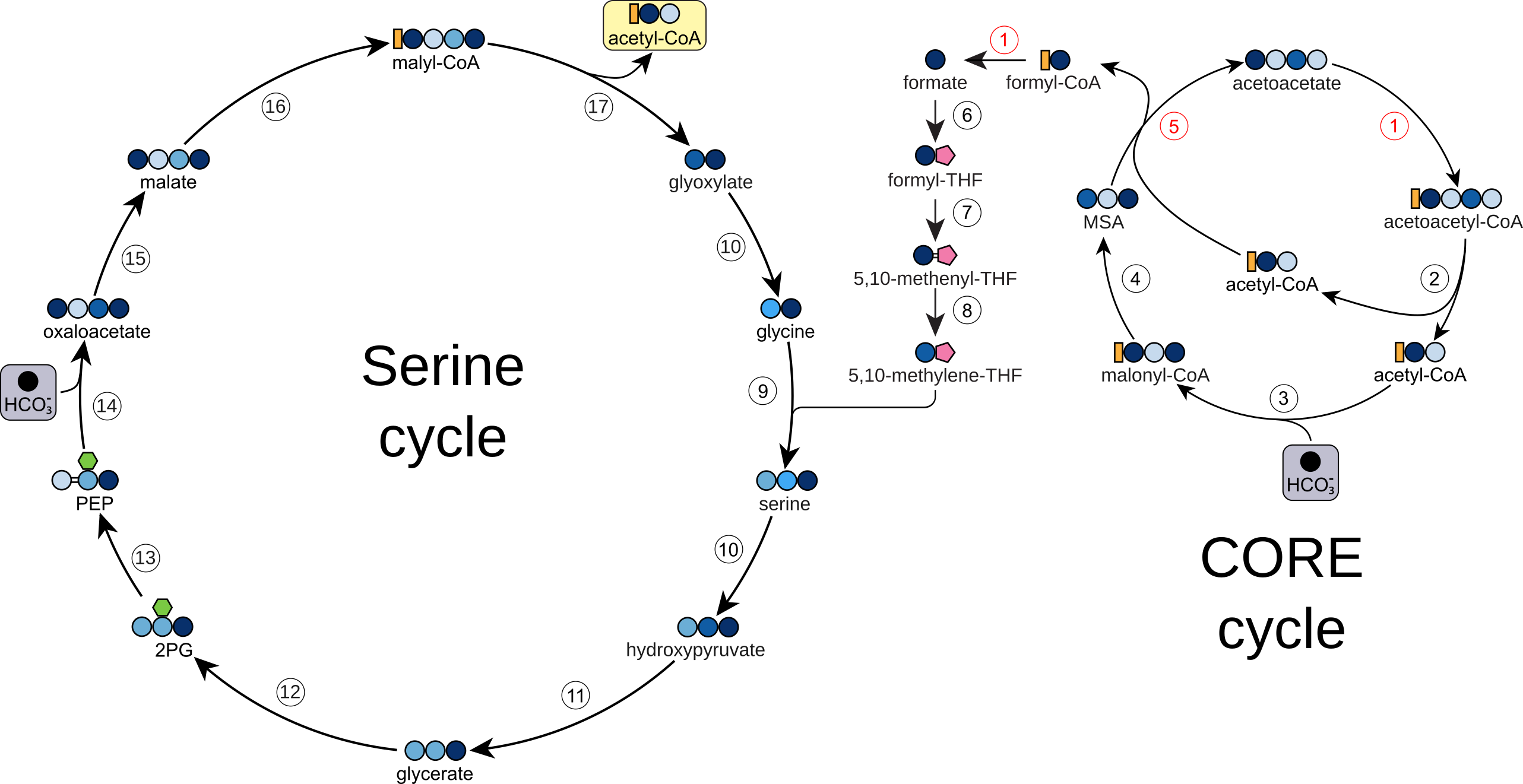

**Figure S9:** The CORE+Serine cycle. This cycle couples CO_2_ reduction to formate via the CORE cycle to formatotrophic growth via the naturally occurring Serine cycle. 1, formyl-CoA:acetoacetate CoA transferase; 2, acetoacetyl-CoA thiolase; 3, acetyl-CoA carboxylase; 4, malonyl-CoA reductase; 5, β-keto-acid cleavage enzyme; 6, formate:tetrahydrofolate ligase; 7, methenyl-tetrahydrofolate ciclohydrolase; 8, methylene-THF dehydrogenase; 9, serine hydroxymethyltransferase; 10, serine:glyoxylate aminotransferase; 11, hydroxypyruvate reductase; 12, glycerate kinase; 13, enolase; 14, PEP carboxylase; 15, malate dehydrogenase; 16, malate thiokinase; 17, malyl-CoA lyase; 2PG, 2-phosphoglycerate; CoA, coenzyme A; MSA, malonate semialdehyde; PEP, phosphoenolpyruvate; THF, tetrahydrofolate.

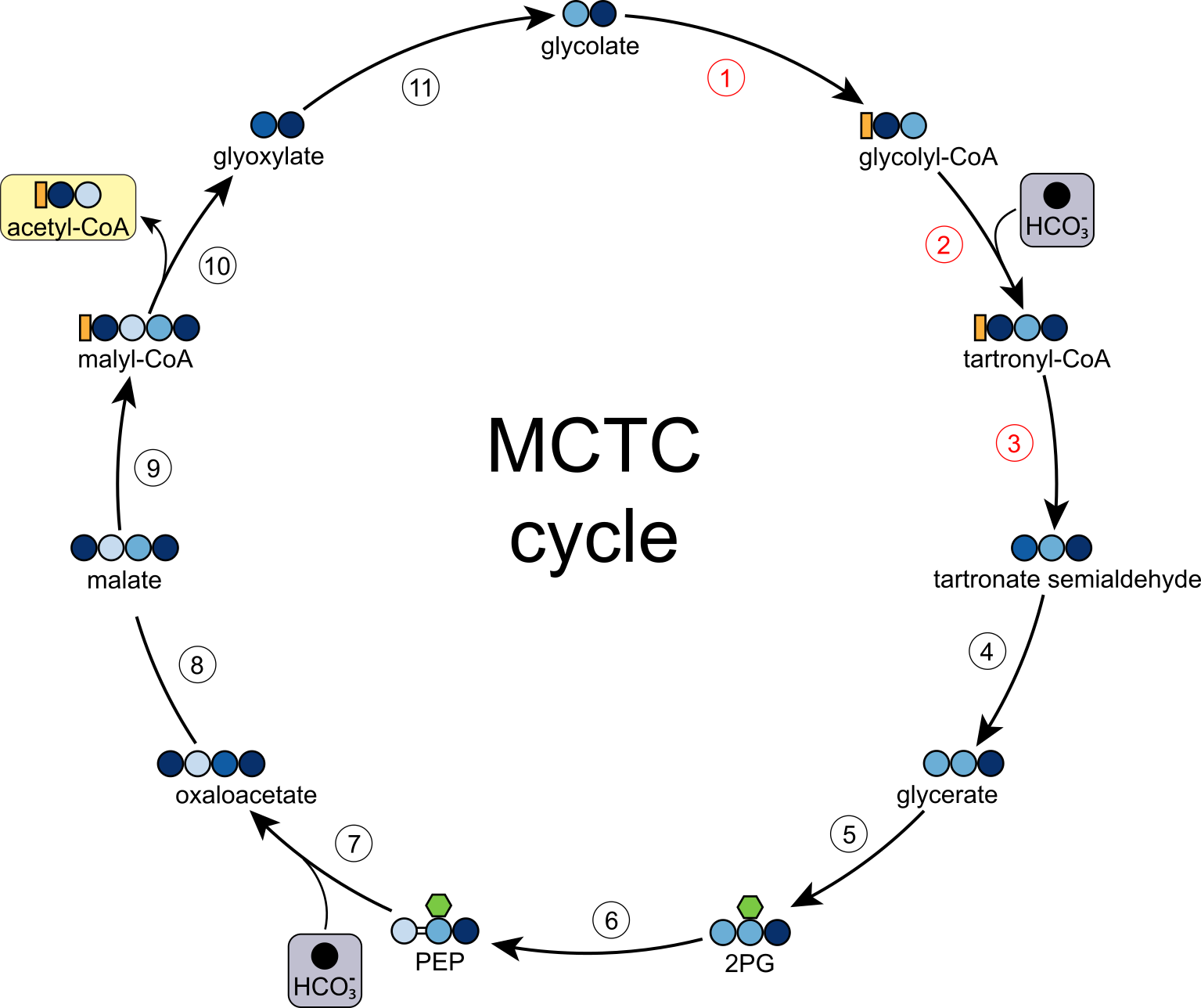

**Figure S10:** The Malyl-CoA-Tartronyl-CoA (MCTC). This cycle is based on the recently reported engineered glycolyl-CoA carboxylase, which is based on a natural biotin-dependent propionyl-CoA carboxylase engineered to accept the smaller substrate glycolyl-CoA^6^. The resulting enzyme has been named glycolyl-CoA carboxylase (GCC). This carboxylation reaction produces tartronyl-CoA, which is further reduced by an engineered malonyl-CoA reductase to tartronate semialdehyde, which is itself reduced to glycerate, a compound that can easily be assimilated into central metabolism. The resulting pathway, known as the tartronyl-CoA (TaCo) pathway, was conceived as a photorespiration bypass, taking 2-phosphoglycolate derived from the Rubisco oxygenation side reaction and recycling it to 3-phosphoglycerate while not only preventing loss of carbon but fixing carbon instead. While this approach is promising, it is currently hampered by the inefficiency of GCC, which hydrolyzes more than 5 ATP per carboxylation reaction, as opposed to 1 ATP for PCC and similar biotin-dependent enzymes. This could be attributed to the difference between propionyl-CoA and glycolyl-CoA as substrates: the latter is not only smaller but contains a hydroxyl group. The presence of a hydroxyl group could destabilize the enolate intermediate required for carboxylation, leading to futile cycles where CO_2_ is released into the active site but is unable to react with the substrate. In a subsequent work, the enzyme was further engineered by using a machine-learning guided approach, obtaining improved variants with reduced futile ATP hydrolysis, albeit at the expense of lower overall catalytic activity^7^. Whether this issue can be fully solved remains an open question. We reasoned that the TaCo pathway could also serve as the basis for a synthetic CO_2_ fixation cycle (Figure 8) where the product, glycerate, goes through lower glycolysis and anaplerosis to malate, which is activated to malyl-CoA and split to acetyl-CoA, which is the end product of this cycle, and glyoxylate, which can be reduced to glycolate to restart the cycle. This pathway has the advantage of being relatively compact, requiring only eleven genes for its implementation, all of which have been tested in vitro, potentially making it easier to engineer, especially in less genetically accessible organisms. Assuming an improved GCC can be engineered, we calculated that this cycle is about 20% more efficient than the CBB cycle at ambient CO_2_, with comparable pathway activity (Table S1, Figure 8). We note that while writing this manuscript, this pathway has also been proposed in another work, in which it has been named the Linear Autocatalytic cycle Tailored for Carbon Harvesting (LATCH)^8^. 1, glycolate-CoA ligase; 2, glycolyl-CoA carboxylase; 3, tartronyl-CoA reductase; 4, tartronate semialdehyde reductase; 5, glycerate kinase; 6, enolase; 7, PEP carboxylase; 8, malate dehydrogenase; 9, malate CoA transferase; 10, malyl-CoA lyase; 11, glyoxylate reductase. 2PG, 2-phosphoglycerate; CoA, coenzyme A; PEP, phosphoenolpyruvate.

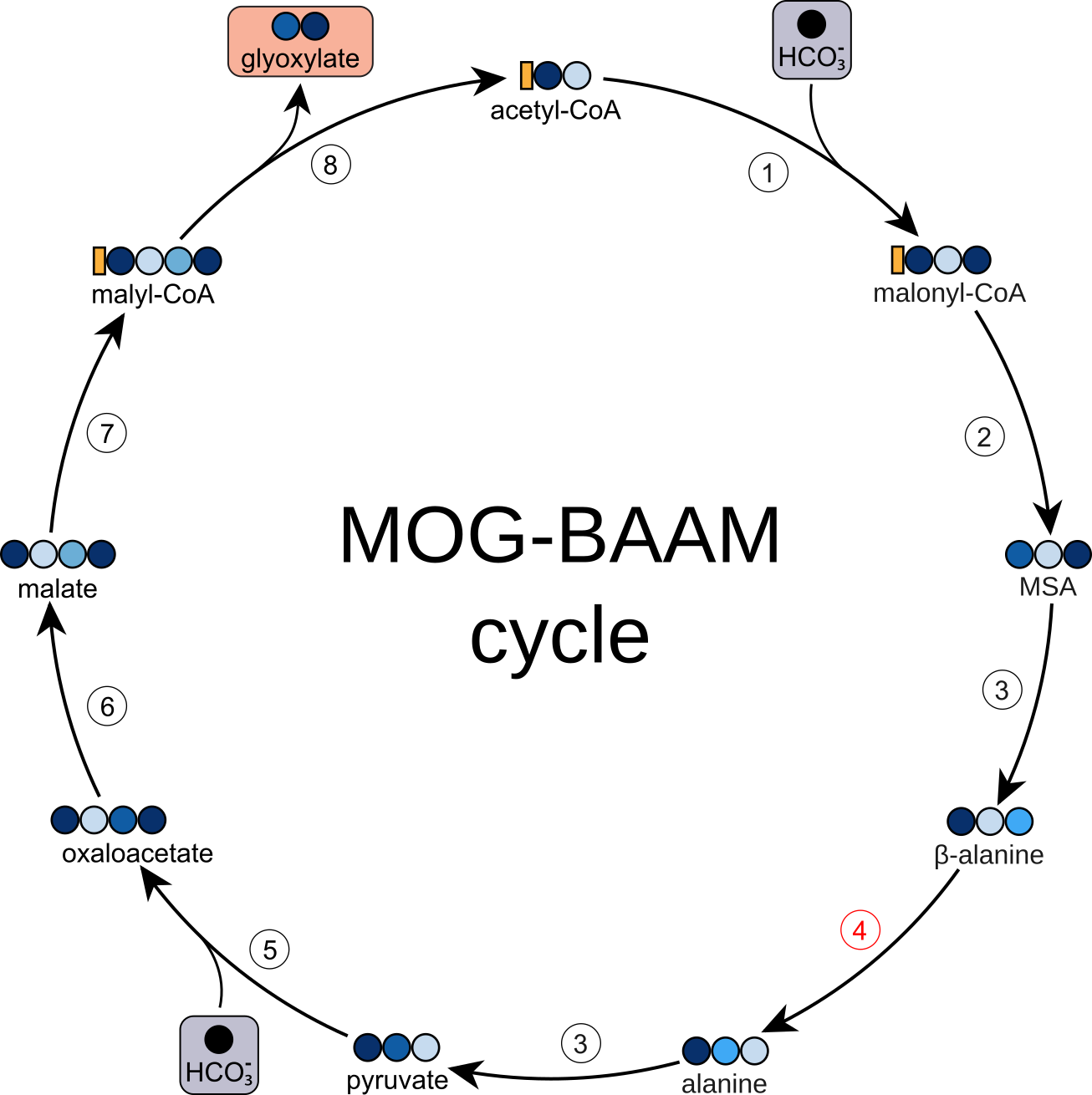

**Figure S11:** The malonyl-CoA-oxaloacetate-glyoxylate-β-alanine aminomutase (MOG-BAAM) cycle^9^. Much like the β-glutamate cycle proposed in this work, this cycle is based on an aminomutase to move the C3 amino group in β-alanine (produced from the transamination of malonate semialdehyde) to C2, yielding alanine, which can be further transaminated to pyruvate. Similar considerations apply to this enzyme as to glutamate aminomutase, with the additional caveat that there is no known alanine aminomutase reported in the literature, not even oxygen-sensitive ones. 1, acetyl-CoA carboxylase; 2, malonyl-CoA reductase; 3, β-alanine:pyruvate aminotransferase; 4, β-alanine aminomutase; 5, pyruvate carboxylase; 6, malate dehydrogenase; 7, malate thiokinase; 8, malyl-CoA lyase; CoA, coenzyme A; MSA, malonate semialdehyde.

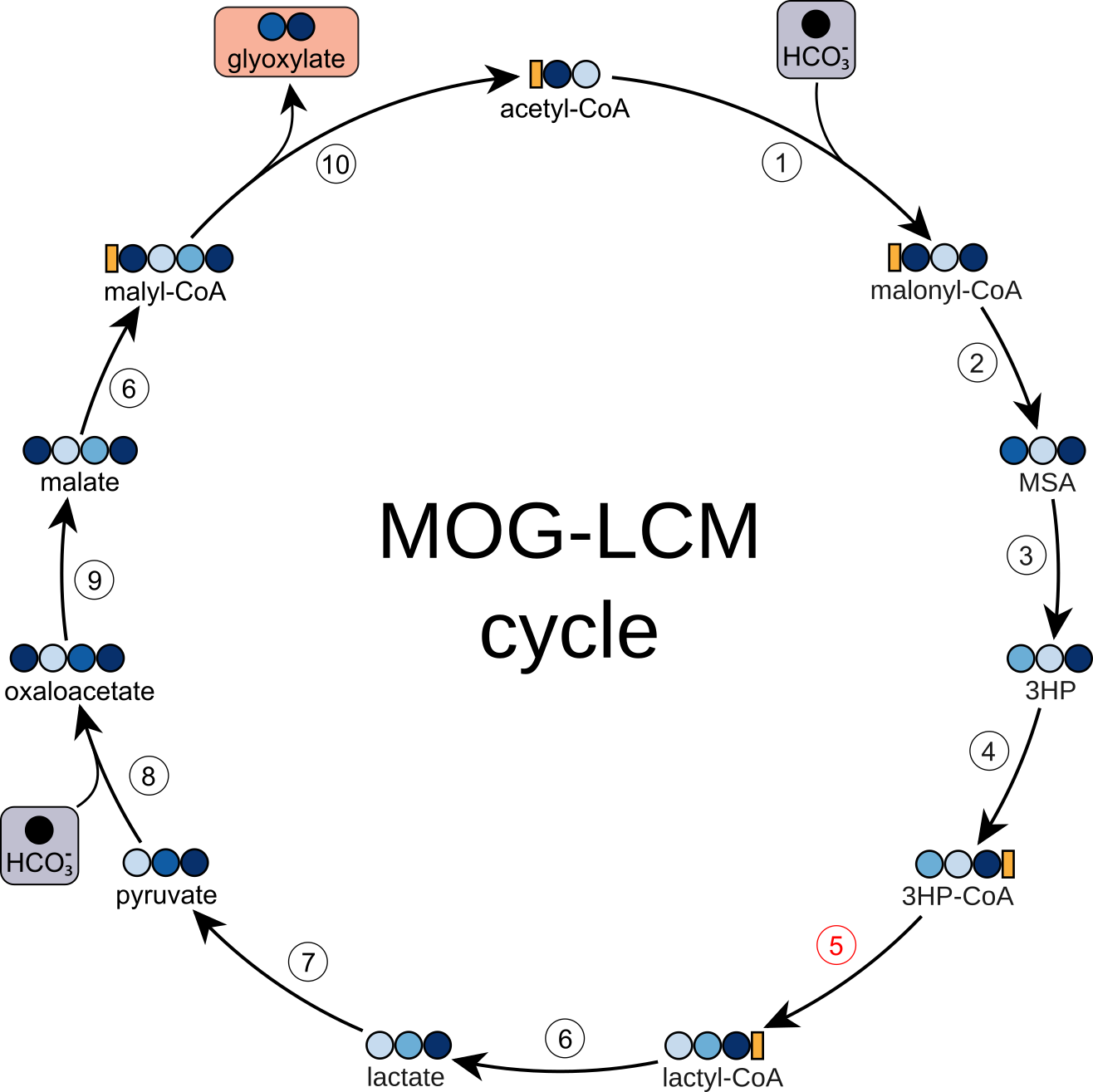

**Figure S12:** The malonyl-CoA-oxaloacetate-glyoxylate-lactyl-CoA mutase (MOG-LCM) cycle^9,10^. This cycle also requires a non-natural isomerase activity, namely lactyl-CoA mutase, converting 3-hydroxypropionyl-CoA to lactyl-CoA. This activity was recently engineered starting from a naturally occurring 3-hydroxybutyryl-CoA mutase^10^. This engineering effort produced an enzyme with a k_cat_ of 0.03, which is around an order of magnitude lower than the activity on its original substrate. We have modeled this cycle using the optimistic scenario where the engineered LCM enzyme can be improved to reach the same k_cat_ as with its native substrate 3-hydroxybutyryl-CoA^1,2^. Even this native activity is rather low, and it is unclear whether it can be improved, making this enzyme a bottleneck for overall pathway activity. Additionally, LCM requires B_12_ as a cofactor, making this cycle unsuitable for implementation in plants and most algae. 1, acetyl-CoA carboxylase; 2, malonyl-CoA reductase; 3, malonate semialdehyde reductase; 4, 3-hydroxypropionate-CoA ligase; 5, lactyl-CoA mutase; 6, lactyl-CoA:malate CoA transferase; 7, lactate dehydrogenase; 8, pyruvate carboxylase; 9, malate dehydrogenase; 10, malyl-CoA lyase; 3HP, 3-hydroxypropionate; 3HP-CoA, 3-hydroxypropionyl-CoA; CoA, coenzyme A; MSA, malonate semialdehyde.

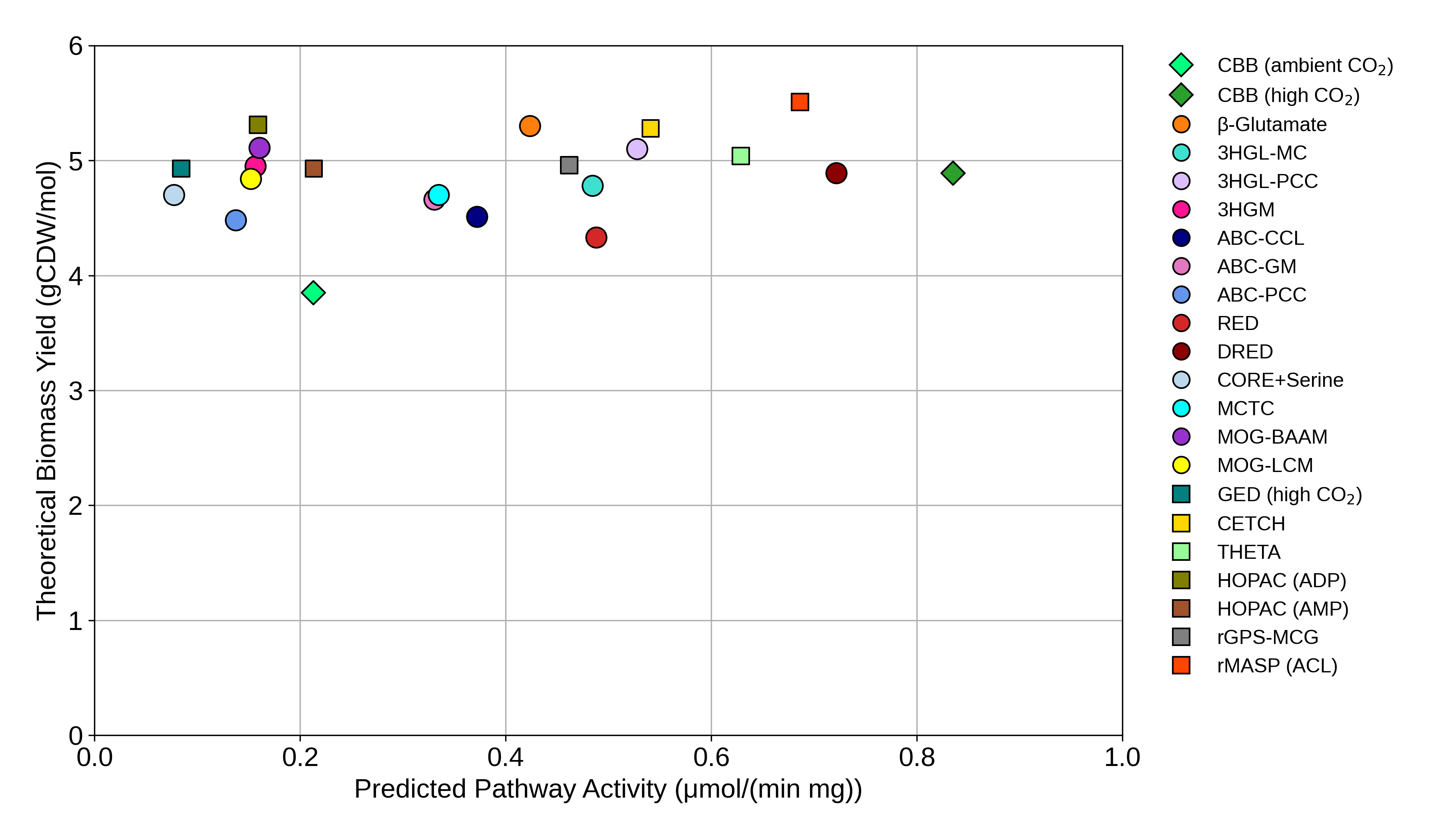

**Figure S13:** Plot of theoretical biomass yield vs predicted pathway activity for each pathway presented in this paper and previously proposed pathways^11–14^. Circles indicate pathways that require enzyme engineering for their implementation, diamonds represent natural pathways, squares represent mix-and-match pathways.

Additional citations from table S2 can be found here^6,10,14–30^.

**References**

1.	Rohwerder, T., Rohde, M.-T., Jehmlich, N. & Purswani, J. Actinobacterial Degradation of 2-Hydroxyisobutyric Acid Proceeds via Acetone and Formyl-CoA by Employing a Thiamine-Dependent Lyase Reaction. *Front. Microbiol.* **11**, 691 (2020).

2.	Kurteva-Yaneva, N. *et al.* Structural Basis of the Stereospecificity of Bacterial B12-dependent 2-Hydroxyisobutyryl-CoA Mutase. *Journal of Biological Chemistry* **290**, 9727–9737 (2015).

3.	Sasikaran, J., Ziemski, M., Zadora, P. K., Fleig, A. & Berg, I. A. Bacterial itaconate degradation promotes pathogenicity. *Nat Chem Biol* **10**, 371–377 (2014).

4.	Roymoulik, I., Moon, N., Dunham, W. R., Ballou, D. P. & Marsh, E. N. G. Rearrangement of L-2-Hydroxyglutarate to L-*threo*-3-Methylmalate Catalyzed by Adenosylcobalamin-Dependent Glutamate Mutase. *Biochemistry* **39**, 10340–10346 (2000).

5.	Pramanik, J. & Keasling, J. D. Effect of Escherichia coli biomass composition on central metabolic fluxes predicted by a stoichiometric model. *Biotechnol. Bioeng.* **60**, 230–238 (1998).

6.	Scheffen, M. *et al.* A new-to-nature carboxylation module to improve natural and synthetic CO2 fixation. *Nature Catalysis* **4**, 105–115 (2021).

7.	Marchal, D. G. *et al.* Machine Learning-Supported Enzyme Engineering toward Improved CO2-Fixation of Glycolyl-CoA Carboxylase. *ACS Synth. Biol.* **12**, 3521–3530 (2023).

8.	Yang, X., Ma, Y., Yan, J., Zhao, G. & Zhang, Y. LATCH: a Linear Autocatalytic cycle Tailored for Carbon Harvesting. *Green Carbon* https://doi.org/10.1016/j.greenca.2025.10.003 (2025) doi:10.1016/j.greenca.2025.10.003.

9.	Bar-Even, A., Noor, E., Lewis, N. E. & Milo, R. Design and analysis of synthetic carbon fixation pathways. *Proceedings of the National Academy of Sciences of the United States of America* **107**, 8889–8894 (2010).

10.	Schulz-Mirbach, H. *et al.* New-to-nature CO2-dependent acetyl-CoA assimilation enabled by an engineered B12-dependent acyl-CoA mutase. *Nat Commun* **15**, 10235 (2024).

11.	Schwander, T., Von Borzyskowski, L. S., Burgener, S., Cortina, N. S. & Erb, T. J. A synthetic pathway for the fixation of carbon dioxide in vitro. *Science* **354**, 900–904 (2016).

12.	Luo, S. *et al.* Construction and modular implementation of the THETA cycle for synthetic CO2 fixation. *Nat Catal* **6**, 1228–1240 (2023).

13.	McLean, R. *et al.* Exploring alternative pathways for the in vitro establishment of the HOPAC cycle for synthetic CO _2_ fixation. *Sci. Adv.* **9**, eadh4299 (2023).

14.	Satanowski, A. *et al.* Awakening a latent carbon fixation cycle in Escherichia coli. *Nature Communications* **11**, 1–14 (2020).

15.	Buckel, W., Dorn, U. & Semmler, R. Glutaconate CoA‐Transferase from *Acidaminococcus fermentans*. *European Journal of Biochemistry* **118**, 315–321 (1981).

16.	Suzuki, S., Yamaguchi, J. & Tokushige, M. Studies on aspartase. I. Purification and molecular properties of aspartase from *Escherichia coli*. *Biochimica et Biophysica Acta (BBA) - Enzymology* **321**, 369–381 (1973).

17.	Ruzicka, F. J. & Frey, P. A. Glutamate 2,3-aminomutase: A new member of the radical SAM superfamily of enzymes. *Biochimica et Biophysica Acta (BBA) - Proteins and Proteomics* **1774**, 286–296 (2007).

18.	Fall, R. *3-Methylcrotonyl-CoA and Geranyl-CoA Carboxylases from Pseudomonas Citronellolis*. (Academic Press, New York London Toronton, 1981). doi:https://doi.org/10.1016/0076-6879(81)71093-0.

19.	Mack, M. *et al.* Biochemical characterization of human 3-methylglutaconyl-CoA hydratase and its role in leucine metabolism. (2006).

20.	Kramer, P. R. & Miziorko, H. M. Purification and characterization of avian liver 3-hydroxy-3-methylglutaryl coenzyme A lyase. *Journal of Biological Chemistry* **255**, 11023–11028 (1980).

21.	Jitrapakdee, S. *et al.* Differential regulation of the yeast isozymes of pyruvate carboxylase and the locus of action of acetyl CoA. *The International Journal of Biochemistry & Cell Biology* **39**, 1211–1223 (2007).

22.	Ferenci, T., Strøm, T. & Quayle, J. R. Purification and properties of 3-hexulose phosphate synthase and phospho-3-hexuloisomerase from *Methylococcus capsulatus*. *Biochemical Journal* **144**, 477–486 (1974).

23.	Fong, S., Machajewski, T. D., Mak, C. C. & Wong, C.-H. Directed evolution of D-2-keto-3-deoxy-6-phosphogluconate aldolase to new variants for the efficient synthesis of D- and L-sugars. *Chemistry & Biology* **7**, 873–883 (2000).

24.	Yin, Y. & Kirsch, J. F. Identification of functional paralog shift mutations: Conversion of *Escherichia coli* malate dehydrogenase to a lactate dehydrogenase. *Proc. Natl. Acad. Sci. U.S.A.* **104**, 17353–17357 (2007).

25.	Muslin, E. H. *et al.* Engineering a domain-locking disulfide into a bacterial malate dehydrogenase produces a redox-sensitive enzyme. *Biophysical Journal* **68**, 2218–2223 (1995).

26.	Toyota, C. G. *et al.* Differential Substrate Specificity and Kinetic Behavior of *Escherichia coli* YfdW and *Oxalobacter formigenes* Formyl Coenzyme A Transferase. *J Bacteriol* **190**, 2556–2564 (2008).

27.	Hügler, M., Menendez, C., Schägger, H. & Fuchs, G. Malonyl-Coenzyme A Reductase from *Chloroflexus aurantiacus* , a Key Enzyme of the 3-Hydroxypropionate Cycle for Autotrophic CO_2_ Fixation. *J Bacteriol* **184**, 2404–2410 (2002).

28.	Chen, D., Ruzicka, F. J. & Frey, P. A. A novel lysine 2,3-aminomutase encoded by the yodO gene of Bacillus subtilis : characterization and the observation of organic radical intermediates. 11 (2000).

29.	Raj, H. *et al.* Characterization of a thermostable methylaspartate ammonia lyase from Carboxydothermus hydrogenoformans. *Appl Microbiol Biotechnol* **94**, 385–397 (2012).

30.	Alban, C., Baldet, P., Axiotis, S. & Douce, R. Purification and Characterization of 3-Methylcrotonyl-Coenzyme A Carboxylase from Higher Plant Mitochondria. *Plant Physiol.* **102**, 957–965 (1993).
